# The prolactin receptor scaffolds Janus kinase 2 via co-structure formation with phosphoinositide-4,5-bisphosphate

**DOI:** 10.1101/2022.11.23.517650

**Authors:** Raul Araya-Secchi, Katrine Bugge, Pernille Seiffert, Amalie Petry, Gitte W. Haxholm, Kresten Lindorff-Larsen, Stine F. Pedersen, Lise Arleth, Birthe B. Kragelund

**Affiliations:** Structural Biophysics, Section for Neutron and X-ray Science, Niels Bohr Institute, University of Copenhagen, 2100 Copenhagen, Denmark; Facultad de Ingenieria Arquitectura y Diseño, Universidad San Sebastian, Bellavista 7, Santiago, Chile; Structural Biology and NMR Laboratory (SBiNLab), Department of Biology, University of Copenhagen, 2200 Copenhagen, Denmark; Section for Cell Biology and Physiology, Department of Biology, University of Copenhagen, 2200 Copenhagen N, Denmark

**Keywords:** IDP, NMR, simulation, prolactin receptor, JAK2, single-pass transmembrane receptors, PI(4, 5)P_2_

## Abstract

Class 1 cytokine receptors transmit signals through the membrane by a single transmembrane helix to an intrinsically disordered cytoplasmic domain that lacks kinase activity. While specific binding to phosphoinositides has been reported for the prolactin receptor (PRLR), the role of lipids in PRLR signalling is unclear. Using an integrative approach combining NMR spectroscopy, cellular signalling experiments, computational modelling and simulation, we demonstrate co-structure formation of the disordered intracellular domain of the human PRLR, the membrane constituent phosphoinositide-4,5-bisphosphate (PI(4,5)P_2_) and the FERM-SH2 domain of the Janus kinase 2 (JAK2). We find that the complex leads to accumulation of PI(4,5)P_2_ at the transmembrane helix interface and that mutation of residues identified to interact specifically with PI(4,5)P_2_ negatively affects PRLR-mediated activation of signal transducer and activator of transcription 5 (STAT5). Facilitated by co-structure formation, the membrane-proximal disordered region arranges into an extended structure. We suggest that the co-structure formed between PRLR, JAK2 and PI(4,5)P_2_ locks the juxtamembrane disordered domain of the PRLR in an extended structure, enabling signal relay from the extracellular to the intracellular domain upon ligand binding. We find that the co-structure exists in different states which we speculate could be relevant for turning signalling on and off. Similar co-structures may be relevant for other non-receptor tyrosine kinases and their receptors.

## Introduction

Cytokine receptors are transmembrane glycoproteins that bind cytokines on the cell surface and transduce signals across the membrane to the interior of the cell. This, in turn, activates signalling pathways leading to multiple outcomes including induction of immune responses, cell proliferation, altered metabolism and differentiation (Brooks et al., 2016). Class 1 cytokine receptors constitute a subclass of receptors that transverse the membrane by a single α-helical transmembrane domain (TMD) (Brooks et al., 2016), separating a folded extracellular domain (ECD) from a disordered intracellular domain (ICD). The prolactin (PRL) receptor (PRLR) is one of the structurally most simple cytokine receptors (***Figure 1***). Signaling by the PRLR/PRL complex is implicated in the regulation of more than 300 biological functions in vertebrates (Bole-Feysot et al., 1998), and its function is especially well-known for its essential role in mammary gland development and lactation (Hannan et al., 2022). Apart from this, deregulation of PRLR/PRL signalling is associated with several pathologies in humans of which hyperprolactinemia resulting in reproductive failure is best described (Bachelot and Binart, 2007; Newey et al., 2013). Deregulation of PRLR/PRL signaling is linked to other diseases and, although still debated, suggested to be implicated in the development and progression of prostate (Sackmann-Sala and Goffin, 2015) and breast (Clevenger and Rui, 2022) cancers.

**Fig. 1:**
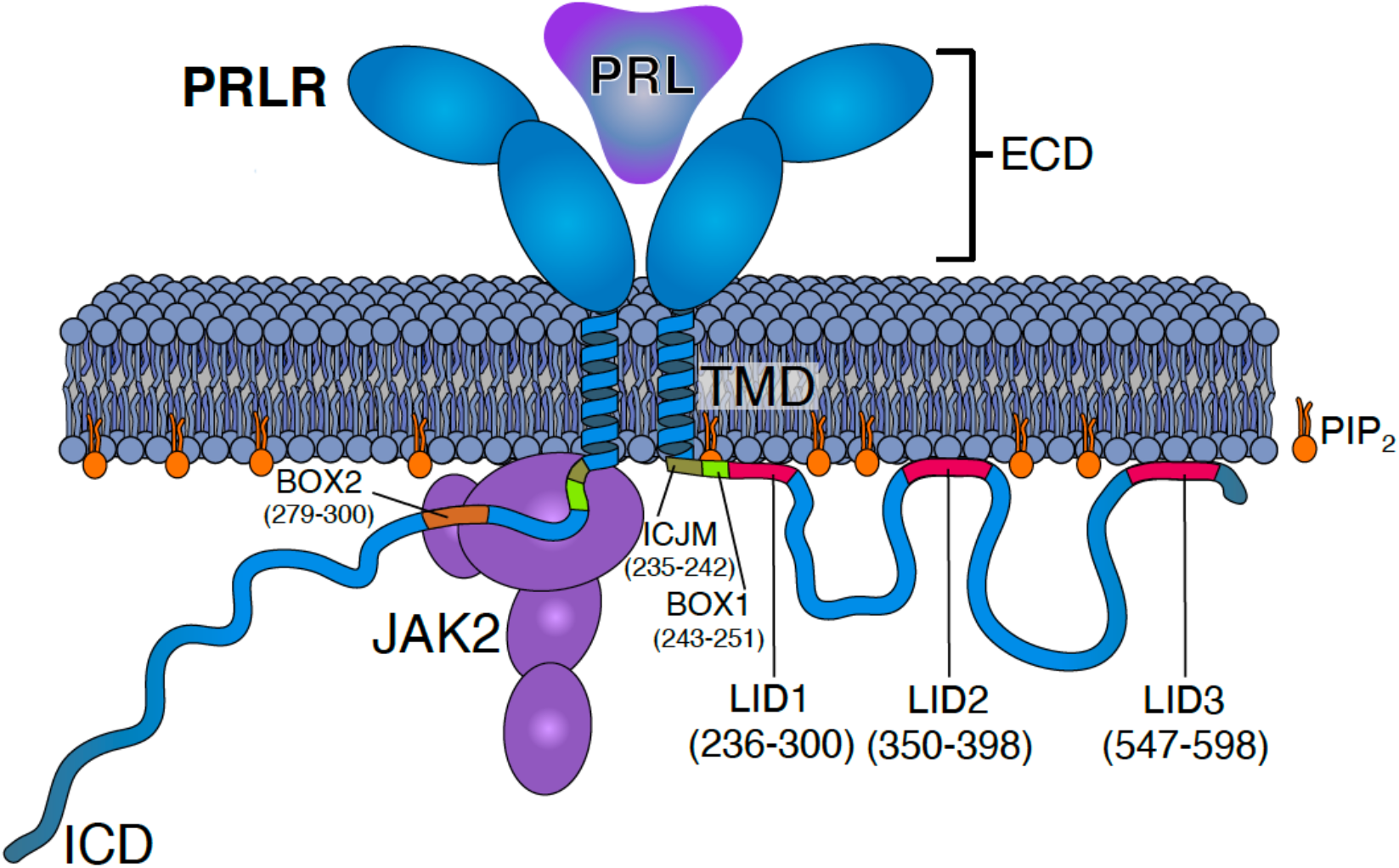
Schematics of the PRLR:PRL:JAK2 complex in the membrane. The PRLR is shown in light blue, the PRL as a dark blue triangle, the PRLR-ICD as a disordered chain and JAK2 in purple. The PI(4,5)P_2_ lipid (PIP2) is shown in orange. The intracellular juxtamembrane (ICJM) region and BOX1 of PRLR-ICD are highlighted in green nuances, while the three LIDs as defined in Haxholm et al., (Haxholm et al., 2015) are highlighted in red. For simplicity only one of the two ICDs is shown associated with JAK2 via the BOX1 (green) and BOX2 (orange) motifs.

For cytokine receptors, signal transduction through the membrane is received by an ICD, which is intrinsically disordered and lacks kinase activity (Haxholm et al., 2015). Thus, association of auxiliary kinases is mandatory for signalling, with the Janus kinases (JAK1–3) and the tyrosine kinase TYK2 being the most thoroughly described (Brooks et al., 2016; Morris et al., 2018). A proline-rich region constituting the so-called BOX1 motif close to the membrane, as well as a second hydrophobic motif termed BOX2, are known to be essential for JAK binding (***Figure 1***) (Ferrao et al., 2018; Rowlinson et al., 2008). However, although progress has been made in the molecular understanding of cytokine binding and despite several structures of folded ECDs (Broutin et al., 2010; de Vos et al., 1992), TMDs (Bocharov et al., 2018; Bugge et al., 2016), a complete receptor (Kassem et al., 2021) and a receptor ICD in complex with JAK1 (Glassman et al., 2022) have emerged, it is still not clear how the signal inside the cell is received by the disordered region to elicit and in turn control signal transduction.

A subset of class 1 cytokine receptors form homodimers and trimeric complexes with their ligands, with the main dimerization occurring in the TMDs (Brooks et al., 2014; Brown et al., 2005; Gadd and Clevenger, 2006; Kubatzky et al., 2001; Seubert et al., 2003). This group includes the growth hormone receptor (GHR), the erythropoietin receptor (EPOR), the thrombopoietin receptor (TPOR) and the PRLR, which have become well-established paradigmatic models. Recently, signal transduction by the GHR has been suggested to occur via a rotation of the transmembrane helices within the dimer leading to a subsequent separation of the ICDs (Brooks et al., 2014; Brown et al., 2005). A torque is hereby exerted on the associated JAK2s, which is thought to relieve inhibition by the pseudokinase domains, initiating signalling. The ICDs of these receptors have been shown to be highly disordered (Haxholm et al., 2015), a feature which is preserved in models of the PRLR (Bugge et al., 2016) and the full-length GHR in nanodiscs (Kassem et al., 2021). This brings forward the question of how signalling is orchestrated by disorder and how a disordered linker region between the TMD and the region bound to the kinases can communicate and transduce information.

For both the PRLR and the GHR, lipid interaction domains (LID) with affinity for negatively charged lipids have been identified in their ICDs (Haxholm et al., 2015). Common to both receptors is that they contain a LID proximal to the membrane, directly overlapping with the JAK2 interaction sites, BOX1 and BOX2 (Seiffert et al., 2020). Using nuclear magnetic resonance (NMR) spectroscopy we identified three LIDs in the PRLR-ICD termed LID1, LID2 and LID3 (***Figure 1***) (Haxholm et al., 2015) and using lipid dot-blots we showed that the PRLR-ICD has variable affinities for different membrane constituents, including for different phosphoinositides (PIs). In particular, PRLR has a distinct affinity for phosphoinositide-4,5-bisphosphate (PI(4,5)P_2_) and lacks affinity for PI(3,4,5)P_2_ (Haxholm et al., 2015). PIs are important constituents of the membrane and play key roles in signalling, both as membrane interaction partners that can be specifically modulated by phosphorylation (Carracedo and Pandolfi, 2008), and as secondary messengers (McLaughlin et al., 2002; Suh and Hille, 2005). Indeed, some single-pass receptors contain conserved anionic lipid interaction sites close to the membrane (Hedger et al., 2015a) and increasing evidence suggest lipid interaction to take on important regulatory roles (McLaughlin et al., 2005; Metcalf et al., 2010). Recently, the epidermal growth factor receptor (EGFR) was shown to sequester PI(4,5)P_2_ by accumulating it around the TMD regulating the dimer/monomer equilibrium and with a possible positive feedback loop through the activation of the phospholipase C (PLC) – diacylglycerol (DAG)-IP_3_ pathways (Maeda et al., 2018). This will lead to subsequent conversion of PI(4,5)P_2_ to PI(3,4,5)P_3_ and hence depletion of PI(4,5)P_2_ from the membrane. Similar depletion of PI(4,5)P_2_ from the plasma membrane has been noted under hypoxia (Lu et al., 2022). For class 1 cytokine receptors the role of PIs in signalling is less clear.

In a cellular context, signalling-related proteins can be membrane anchored through modifications such as acylation (Patwardhan and Resh, 2010; Rawat et al., 2013; Rawat and Nagaraj, 2010) or via designated lipid-binding domains. This includes the four point-1, ezrin, radixin moesin (FERM) domain of radixin, focal adhesion kinase (FAK) and the protein tyrosine phosphatase L1 (PTPL1) (Bompard et al., 2003; Feng and Mertz, 2015; Hamada et al., 2000), the SH2 domains of the Src family kinases (Park et al., 2016; Sheng et al., 2016) and the FERM-SH2 domain of merlin (Mani et al., 2011). Thus, kinases and receptors can co-localize at the plasma membrane without necessarily being bound within a complex. It is, however, unclear whether such membrane co-localisation has functional consequences, such as enhancing signalling speed, and whether the membrane may act as an additional scaffolding platform that enhances binding via restriction in the two-dimensional plane.

Recent work on disorder in membrane proteins and on the interplay between membrane proteins and lipids have revealed the need for strong integrative methods, combining successfully various structural biology techniques, biophysics and computational biology (Basak et al., 2022; Larsen et al., 2022). These include NMR, small-angle X-ray scattering, crosslinking-mass spectrometry and single molecule fluorescence combined with molecular dynamics simulations (Chavent et al., 2018; Goretzki et al., 2023). These efforts have provided important insights into the role of lipids in regulation of membrane proteins. For TRPV4, a member of the TRP vanilloid channel family, it was shown that an autoinhibitory patch of the receptor competed with PI(4,5)P_2_ binding at the membrane to attenuate channel activity, and MD simulation showed that lipid binding affected the ensemble dynamics(Goretzki et al., 2023). For EphA2, a receptor tyrosine kinase, an integrative study showed how PIs mediate the interaction between the kinase domains, facilitated by clustering of PIPs via binding to the receptor juxtamembrane domain (Chavent et al., 2018) further promoting conformation specific dimerization (Stefanski et al., 2021). Thus, studying dynamic processes at the membrane interface is an emerging field requiring integrative structural biology approaches for detailed atomic resolution information.

For the PRLR it is still not clear whether, and if so how, interactions between the ICD and the membrane impact signal transduction and association with JAK2. Nor is it understood how structural disorder can relay and transmit information from the TMD to initiate signalling. To shed light on the molecular details underlying a potential interplay between the receptor, membrane and kinase we focused on the human PRLR and its LID1 closest to the membrane, facilitating the first intracellular step in signaling. Using an integrative approach combining NMR spectroscopy, cell biology and computational modelling, we demonstrate the formation of a co-structure comprised of the disordered PRLR-ICD, the membrane constituent PI(4,5)P_2_ and the FERM-SH2 domain of the JAK2. Facilitated by this co-structure, the disordered region closest to the membrane forms an extended structure, which we suggest stabilizes the disordered domain, allowing signal relay from the extracellular to the intracellular domain.

## Results

### LID1 is disordered in solution and when tethered to the transmembrane helix

Membrane interactions by PRLR-ICD have previously been studied in the absence of anchoring to the TMD defining three LIDs, with LID1 closest to the membrane (Haxholm et al., 2015). Since tethering would increase the local concentrations at the membrane and the ICD, this could affect affinity, complex lifetime as well as the degree by which structure formation would be captured. Furthermore, the first LID, LID1, is disordered and is located in the juxtamembrane region where it is responsible for transmitting information on extracellular hormone binding to the bound JAK2. As this constitutes the very first step on the intracellular side, and given the distance to the other two LIDs (LID2 and LID3) and their disconnect from the TMD by long disordered regions, we focused exclusively on LID1. We recombinantly expressed the TMD (residues 211–235 with numbering corresponding to the processed protein) with five residues added at the two termini (TMD_F206-V240_), and the TMD with the first 35 residues of LID1, TMD-ICD_F206-S270_ (***Figure 2A***). We then examined their structural propensities in detergents and in small unilamellar vesicles (SUVs) by NMR spectroscopy. In 1,2-dihexanoyl-sn-glycero-3-phosphocholine (DHPC) micelles, most of the TMD resonances of the TMD_F206-V240_ and TMD-ICD_F206-S270_ variants were readily superimposable in the ^15^N,^1^H-HSQC spectra (***Figure 2 - figure supplement 1***), suggesting that the conformation of the TMD was not affected by the presence of the ICD. For the TMD-ICD_F206-S270_, C^α^ NMR resonances were collected for most of the disordered region, while backbone carbon resonances for the TMD, except for A222, and the region G236-P246 immediately following it, were broadened beyond detection in the 3D spectra, preventing assignments (***Figure 2B***). This may suggest that the first ten residues of the ICD interact with or are buried in the micelles, or are affected by the overall slower tumbling of the micelle, whereas the complete overlap of the TMD residues in the ^15^N-HSQC spectra confirm the helical structure as seen previously. We assigned the backbone nuclei of the detectable resonances of TMD-ICD_F206-S270_ in DHPC micelles and compared the secondary chemical shifts (SCS) to those of the ICD alone (ICD_G236-Q396_) (***Figure 2C***). Whereas the region of the ICD that is undetected in TMD-ICD_F206-S270_ formed transient extended structures in the absence of the TMD, the observable parts were highly similar suggesting lack of structure induction by TMD tethering or the micelles. In 1-palmitoyl-2-oleoyl-sn-glycero-3-phosphocholine (POPC) SUVs, only the resonances of the most C-terminal residues of TMD-ICD_F206-S270_ were detectable; however, the chemical shifts suggested that the residues were disordered (***Figure 2 – figure supplement 2***). Taken together, these results suggest that most of the ICD residues remain disordered when tethered to the TMD and in the presence of a neutral membrane mimetic.

**Fig. 2:**
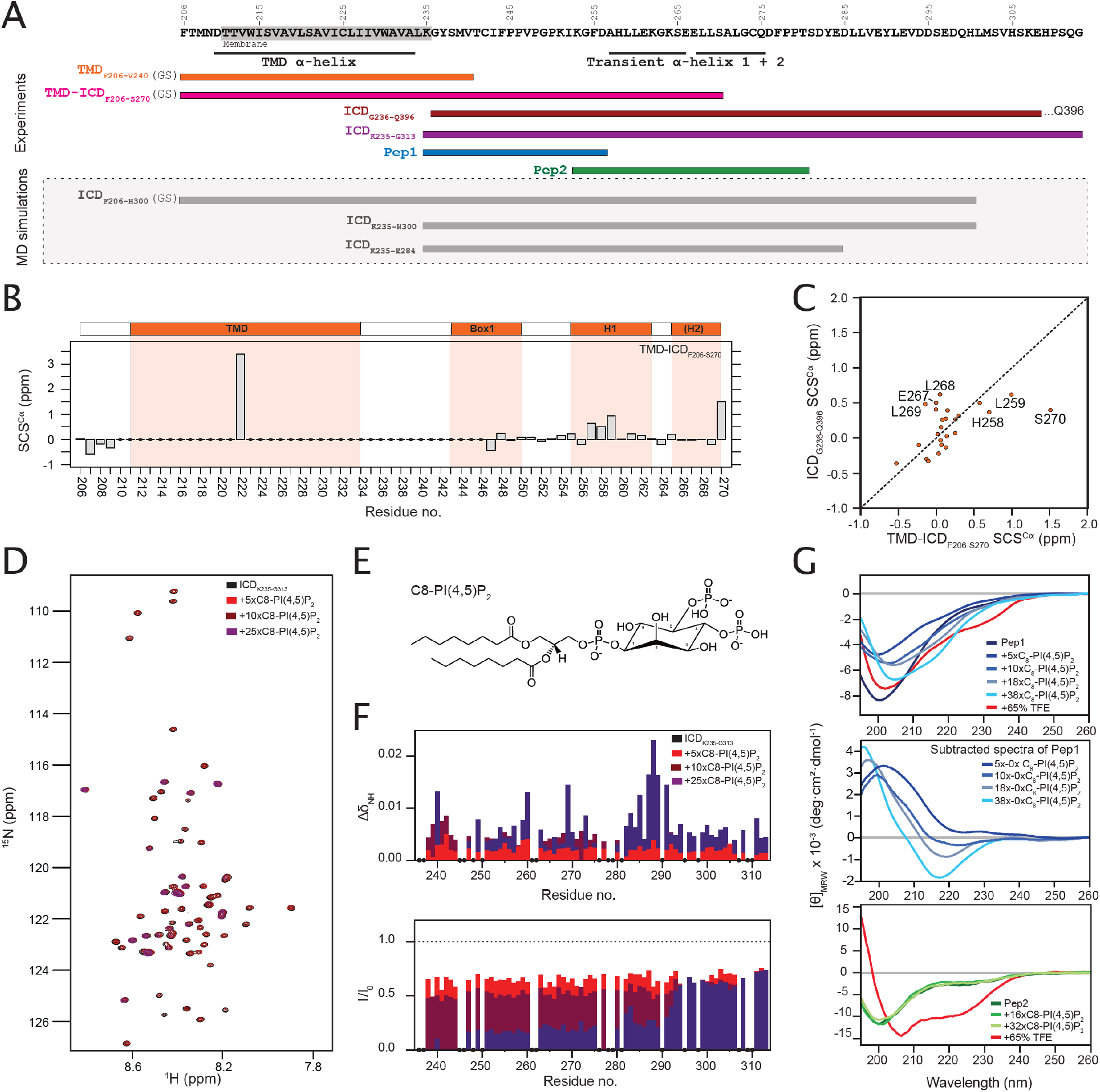
The ICJM region of the PRLR interacts with PI(4,5)P_2_. A) Overview of investigated PRLR variants. **B)** Secondary chemical shifts (SCSs) of TMD-ICD_F206-S270_ reconstituted in DHPC micelles. **C)** Correlation plot of the SCSs of ICD_G236-Q396_ plotted against those of TMD-ICD_F206-S270_. **D)** ^15^N,^1^H-HSQC spectra of ^15^N-ICD_K235-G313_ titrated with 5x, 10x and 25x molar excess of C_8_-PI(4,5)P_2_. **E)** Structure of C_8_-PI(4,5)P_2_. **F)** Backbone amide chemical shift perturbations (CSPs) and peak intensity changes upon addition of C_8_-PI(4,5)P_2_ to ^15^N-ICD_K235-G313_ plotted against residue number. **G) Top**: Far-UV CD spectra of Pep1 titrated with C_8_-PI(4,5)P_2_ or in 65% TFE. **Middle:** Far-UV CD spectra of Pep1 in the presence of 5x-38x C_8_-PI(4,5)P_2_ subtracted with the spectrum of Pep1 in the absence of C_8_-PI(4,5)P_2._ **Bottom**: Far-UV CD spectra of Pep2 titrated with C_8_-PI(4,5)P_2_ or in 65% TFE. * indicate missing data points. **Figure supplement 1**: *^15^N, ^1^H-HSQC spectra of A) TMD_F206-V240_ and TMD-ICD_F206-S270_ in DHPC micelles, and (B) TMD-ICD_F206-S270_ in POPC SUVs.* **Figure supplement 2**: *C^α^ secondary chemical shifts of ICD_G236-Q396_*

### LID1 binds PI(4,5)P_2_ in the juxtamembrane region forming extended structures

PRLR-ICD has previously been shown to bind PI(4,5)P_2_ (but not PI(3,4,5)P_3_) (Haxholm et al., 2015), suggesting that this lipid could modulate membrane affinity and the structural properties of the PRLR-ICD. To separate headgroup effects from lipid bilayer surface effects, we used a short-chain C_8_-PI(4,5)P_2_, which has a high critical micelle concentration (CMC) of 2 mM (Goñi et al., 2014) and analysed the structural changes by NMR and CD spectroscopy at concentrations below the CMC (Goñi et al., 2014) to identify the binding site.

^15^N-labelled ICD_K235-G313_ covering LID1 (***Figure 1***) was titrated with C_8_-PI(4,5)P_2_ and binding was assessed by ^1^H-^15^N-HSQC spectra (***Figure 2D-F***). The chemical shift perturbations (CSPs) were modest whereas substantial intensity changes were observed throughout the chain, supporting a direct interaction between the ICD and lipids. The resonances from G236-F244 completely disappeared suggesting exchange on an intermediate NMR timescale, while intensities were substantially reduced in the V247-S290 region (***Figure 2F***). In the region from D285-E292 we observed an almost inverse correlation between the CSPs and the intensities. This suggests that in contrast to the preceding region, a faster local exchange rate allows us to follow the resonances from the bound state in this region, giving rise to the large CSPs. From this region and to the C-terminus, only moderate intensity changes were observed (***Figure 2F***). These findings suggest that the primary PI(4,5)P_2_ binding site is located closest to the membrane in what we define as the intracellular juxtamembrane (ICJM) region (K235-C242). The ICJM is located N-terminally to the BOX1 motif (_243_IFPPVPGPK_251_ (UNIPROT); _245_PPVPGPK_251_ (elm.eu.org)).

As the resonance-broadening precluded observation of the bound state, two overlapping peptides, Pep1 (K235-D256) and Pep2 (K253-T280), were constructed and evaluated by CD spectroscopy. In isolation, the peptides were disordered as judged by the negative ellipticity at 200 nm in their far-UV CD spectra (***Figure 2G***). Pep1 and Pep2 were titrated with C_8_-PI(4,5)P_2_ and the structural changes monitored (***Figure 2G***). For Pep2, the far-UV CD signal was unaffected by the presence of C_8_-PI(4,5)P_2_. In contrast, for Pep1, large spectral changes were seen, which were unrelated to helix formation. Subtracting the spectra in the presence and absence of C_8_-PI(4,5)P_2_, revealed a negative ellipticity minimum at 218 nm, a strong indicator of β-strands, showing that when bound to C_8_-PI(4,5)P_2_, a distinct extended (strand-like structure) signature was seen (***Figure 2G***). This suggests that this region of LID1 changes its conformational properties upon binding to C_8_-PI(4,5)P_2_. We evaluated the intrinsic helical propensities of the two ICD segments by exposure to high trifluorethanol (TFE) concentrations. Here, Pep2 was readily able to form helical structure as expected from the presence of two transient helices (Haxholm et al., 2015) (***Figure 2G, top***), whereas Pep1 was not (***Figure 2G, bottom***).

In summary, LID1 of the PRLR-ICD interacts with PI(4,5)P_2_,with the primary interaction site located in the K235-S290 region. Headgroup interaction was dominantly located to the region K235-D256 constituting the ICJM and the BOX1 motif and this interaction induced the formation of a regional extended structure in the PRLR-ICD.

### LID1 has specific PI(4,5)P_2_ contacts which drive PI(4,5)P_2_ recruitment

To obtain a more detailed characterization of the behaviour of the disordered PRLR-ICD near lipid bilayers, as well as the effect of different anionic lipid headgroups in these, we turned to molecular simulations. Here, as explained above, we focused on the LID1 (K235-H300), alone and in context of TMD, and first placed a coarse-grained (CG) model of the TMD-ICD_F206-H300_ in three different mixed-membrane systems. These contained an upper leaflet consisting of 100% POPC and lower leaflets composed of a 90:5:5 or 80:10:10 mixture of POPC:POPS:PI(4,5)P_2_ (***Figure 3AB***) or a 70:30 molar ratio mixture of POPC:1-palmitoyl-2-oleoyl-sn-glycero-3-phospho-L-serine (POPS) (***Figure 3 – figure supplement 1EF***). Since the Martini forcefield may produce unrealistically collapsed disordered regions, increasing the strength of the protein-water interactions by 8–10% has provided satisfactory results when applied to the simulation of other disordered regions or multidomain proteins (Thomasen et al., 2022). Thus, the simulations were run using a modified version of the Martini3 forcefield with a 10% increase in the strength of the protein-water interactions. For comparison, similar simulations were performed using the Martini2 forcefield (***Figure 3 – figure supplement 1***).

We first analysed the dynamics of the LID1 during the simulations focusing on the pattern of protein-lipid contacts. Here, we measured the number of protein-lipid contacts focusing either on interactions between the protein and lipid headgroups or the protein and lipid acyl chains. In both cases we determined the fraction of the simulation time that the protein and different parts of the lipid were within 7Å of each other. In general, we observed that residues in the N-terminal part of the LID1 (K235 - D255), which includes the ICJM and BOX1, established contacts with the bilayer in all three membrane systems (***Figure 3A,BFigure 3 – figure supplement 2***) Furthermore, a hydrophobic region rich in prolines (V240–P250) made consistent contacts with the acyl-chains and much more than to the headgroups, indicating penetration into the lower-leaflet. Similar behaviour has been reported with CG-MD simulations for the juxtamembrane region of other single-pass transmembrane receptors (Hedger et al., 2015b). For PRLR, the pattern of interaction was independent on the lipid composition, at least in terms of protein-POPC contacts, and the region interacting with the lipids was similar in all three membrane systems (***Figure 3A,B, Figure 3 – figure supplement 2***).

**Fig. 3.**
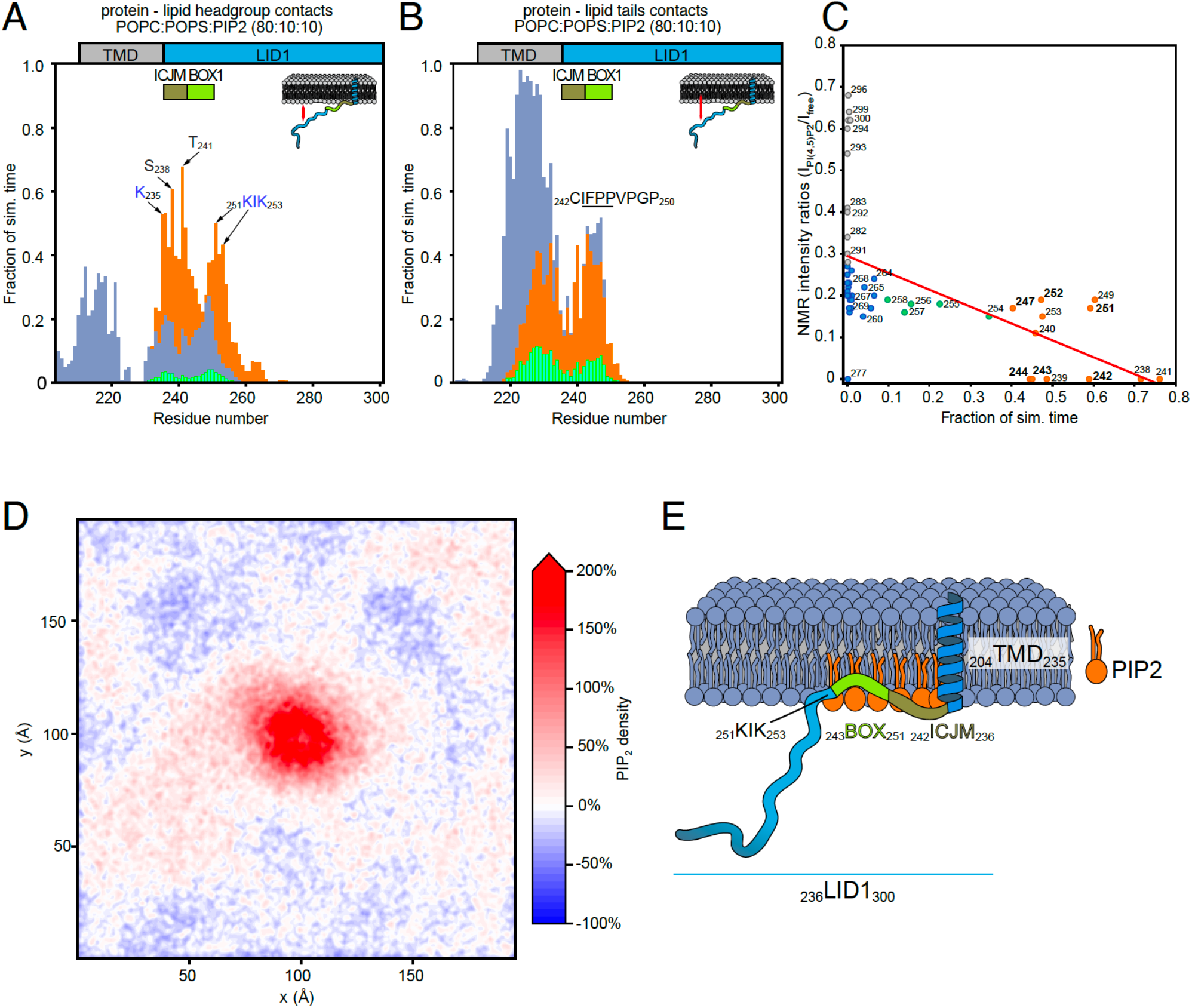
Protein – lipid interactions of PRLR-ICD_LID1_ obtained from CG-MD simulations. (A-B) Protein – lipids contact histograms for PRLR-ICD_LID1_+POPC:POPS:PI(4,5)P_2_ (80:10:10). **A**) Contacts between the protein and lipid headgroups. A contact is counted if the distance between the backbone beads of the protein is ≤ 7 Å from the head-group beads of the lipids. **B**) Contacts between the protein and the acyl chains of the lipids. A contact is counted if the distance between the backbone bead of the protein is ≤ 7 Å from the acyl-chain bead of the lipids. **C**) Correlation between the change in NMR signal and the contact frequency between PRLR-ICD_LID1_ and the lipid headgroups from the PRLR-ICD_LID1_ + POPC:POPS:PI(4,5)P_2_ (80:10:10) system. Pearson correlation coefficient of -0.55 with *p* = 4.0x10^-5^ and *R^2^*= 0.3. **D**) Average PI(4,5)P_2_ density map (*xy*-plane) taken from the PRLR-ICD_LID1_ + POPC:POPS:PI(4,5)P_2_ (80:10:10) simulation. The map is colored according to the enrichment/depletion percentage with respect to the average density value. **E**) Schematic representation of how the interactions and the embedment into the membrane of PRLR contribute to the co-structure formation. The data from the simulations correspond to those of the production stage (see methods). **Figure supplement 1**: *Protein – lipid interactions of PRLR-ICD_LID1_ obtained from CG-MD simulations using the martini 3.0b3.2 forcefield* **Figure supplement 2**: Complementary analysis of the Protein e Protein – lipid interactions of PRLR-ICD_LID1_ obtained from CG-MD simulations.

Although the extent and pattern of protein-lipid interactions appeared constant, a striking observation was made in both systems containing PI(4,5)P_2_. Here, protein-lipid interactions between residues K235 and K253 and PI(4,5)P_2_ were present during a large fraction (250%) of the simulations, despite PI(4,5)P_2_ being present at only 5% or 10% of the total lipids (***Figures 3 – figure supplement 2***). This was also observed in simulations with the Martini2 forcefield, in which the LID1 promptly collapsed in a globular and unstructured coil (***Figure 3 – figure supplement 1***). This suggested that PI(4,5)P_2_ spontaneously accumulated, or in other ways become concentrated around the TMD-ICD_F206-H300_. The computed average density maps for PI(4,5)P_2_ indeed showed that PI(4,5)P_2_ formed a microdomain around the TMD (***Figure 3D***). The low frequency of contacts between the protein and POPS suggests that POPS did not accumulate or compete with PI(4,5)P_2_ for binding to the TMD-ICD_F206-H300_, further supporting the preference for PI(4,5)P_2_ observed earlier (Haxholm et al., 2015). Similar preference was also observed with 5% PI(4,5)P_2_ (***Figure 3 – figure supplement 2***) as well as with Martini2 (***Figure 3 – figure supplement 2E-F***).

A more detailed analysis of LID1-PI(4,5)P_2_ contacts revealed a preference for certain residues, shown as peaks in the protein-headgroup contact profiles. In particular K235, S238, T241, and K251 and K253, which define a KIK motif suggested as a PI(4,5)P_2_ binding motif (Kjaergaard and Kragelund, 2017), engaged in highly populated contacts (***Figure 3A and figure supplement 2A***). The pattern of contacts was not affected by PI(4,5)P_2_ concentration; however, the frequency of contacts almost doubled as a result of the increase from 5% to 10% of PI(4,5)P_2_. The hydrophobic residues in the ICJM and BOX1 penetrate the headgroup layer and facilitate the approximation of the KIK motif to the PI(4,5)P_2_ headgroups. The stabilization of the structure provided by the hydrophobic residues from the ICJM and BOX1 is also reflected in their decrease in flexibility, observed as a shoulder on the RMSF-BB plot, for the residues that comprise the ICJM and BOX1 of PRLR-LID1. Very similar profiles of the RMSF-BB plot was obtained for the systems with respectively 5 and 10% PI(4,5)P_2_ in POPC:POPS, suggesting that the loss in flexibility is coupled to the buried hydrophobic residues rather than to specific PI(4,5)P_2_ interaction (***Figure 3 – figure supplement 2C).*** Contributions from other positively charged residues such as K262 and K264 were very small. To validate the observations from the simulations, we compared the pattern of protein:PI(4,5)P_2_ interactions observed in the NMR experiments to those from the simulations containing PI(4,5)P_2_ (***Figure 3C***). A clear correlation between loss of NMR signal and high frequency of protein-PI(4,5)P_2_ and POPC/POPS contacts in the 80:10:10 simulation was observed, reinforcing that the simulations are able to capture the specificity of protein-PI(4,5)P_2_ interactions. Furthermore, both experiments (***Figure 2***) and simulations (***Figure 3***) show that the residues involved in binding to PI(4,5)P_2_-containing membranes overlap with those that are involved in binding to JAK2.

In summary, the CG-simulations of TMD-ICD_F206-H300_ near lipid membranes showed accumulation of PI(4,5)P_2_ around the TMD and the N-terminus of LID1 involving the ICJM and BOX1. The residues made contact with the membrane independently of lipid type, with BOX1 residues acting as a tether by penetrating the headgroups. This tethering keeps positively charged residues, such as K251 and K253, near the membrane. This may be the driver for PI(4,5)P_2_ recruitment, enhanced by higher PI(4,5)P_2_ concentration. Intriguingly, we observed that the same regions involved in JAK2 binding (BOX1), also play roles in membrane association and lipid recruitment.

### JAK2-FERM-SH2 and PRLR-ICD_LID1_ form co-structures with PI(4,5)P_2_ on membranes

It has been suggested that JAK2 and PRLR interact constitutively in cells (Campbell et al., 1994; Rui et al., 1994) although recent data for the GHR have shown that the Src family kinase Lyn competes for this site (Chhabra et al., 2023). Thus, given our observations that residues from LID1 form lipid-specific contacts with the membrane constituents using the same region covering the binding interface with the FERM domain of JAK2, we decided to explore the structure and dynamics of the JAK2(FERM-SH2):PRLR(LID1) complex near lipid bilayers (***Figure 4***). To do so, an atomistic model of the complex of a smaller region of PRLR-ICD_K235-E284_ bound to the JAK2-FERM-SH2 domains (residues P37 to T514) was built, taking advantage of crystal structures of JAK2-FERM-SH2 and of JAK1-FERM-SH2 and TYK2-FERM-SH2 bound to analogous fragments of the ICDs of the interferon λ- and α-receptors (IFNLR1 and IFNAR1), respectively. This model was used to perform all-atom MD simulations in a water-box to obtain equilibrated structures for further simulations, and to study the dynamics of the protein complex. The average contact map between JAK2-FERM-SH2_P37-T514_ and PRLR-ICD_K235-E284_ showed clusters of contacts in which residues from BOX1 of LID1 formed close contacts (avg. dist. ≤ 4Å) with residues from the F2 lobe (and the F1-F2 linker) and the SH2 domain of JAK2-FERM-SH2, respectively (***Figure 4 – figure supplement 1A***). C-terminally to BOX1, a second set of persistent contacts was observed, again involving charged and hydrophobic residues including F255, L259, E261, K262 and K264 from PRLR-ICD_K235-E284_. Conservation analysis using ConSurf (Ashkenazy et al., 2016; Landau et al., 2005) (***Figure 4 – figure supplement 1B-D***) showed conserved residues in the interface, particularly those of BOX1 of PRLR (P245, P248, K251, I252) (***Figure 4 – figure supplement 1B),*** while a patch of conserved residues (T225, I229 and F236) in JAK2-FERM-SH2 formed close contact with residues from PRLR BOX1. JAK2 residues V183 and L184 interacted with the backbone of _251_KIK_253_ of PRLR-ICD, whereas other, less conserved residues such as E173 and E177 formed transient salt-bridges with K251 and K253 of PRLR-ICD (see ***Figure 4 – figure supplement 1A)***. The contact map also showed that the N-terminus of the ICJM remained flexible without close contacts with JAK2-FERM-SH2 (avg. dist. ≥ 6Å).

**Fig. 4.**
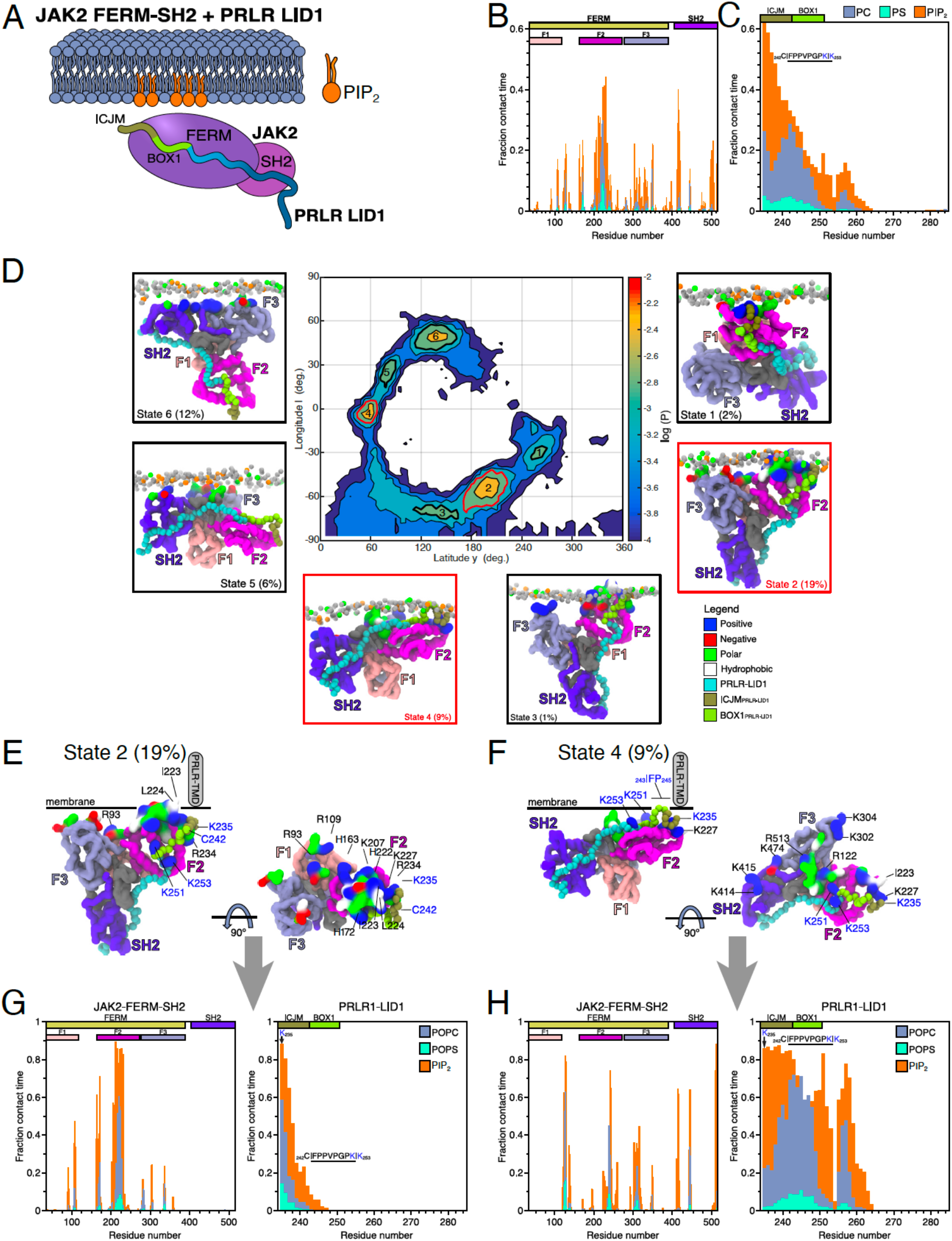
Protein – lipid interactions of the JAK2-FERM-SH2 PRLR-ICD_LID1_ complex obtained from CG-MD simulations. **A)** Schematic representation of the simulated system. Combined **B)** JAK2-FERM-SH2-lipid and **C)** PRLR-ICD_LID1_-lipid contact frequency histograms for the 16 CG simulations of the JAK2-FERM-SH2 +PRLR-ICD_LID1_+ POPC:POPS:PI(4,5)P_2_ system. **D)** Distribution of the orientations adopted by the JAK2-FERM-SH2 + PRLR-ICD_LID1_ complex when bound to lipids taken from the 16 simulations with POPC:POPS:PI(4,5)P_2_ in the lower-leaflet. The snapshots surrounding the map correspond to representative conformations of the highlighted states also indicating the fraction total bound time for which each state was observed. Representative conformations of **E)** State 2 and **F)** State 4. The grey cylinder depicts the position where PRLR-TMD should be located. Representative protein-lipid contact histograms for **G)** State2 and **H)** State4 colored as in panels B and C. **Figure supplement 1**: *Analysis of the JAK2-FERM-SH2-PRLR-ICD_LID1_ AA-MD simulation*. **Figure supplement 2.** *Complementary analysis of Protein – lipid interactions of the JAK2-FERM-SH2 PRLR-ICD_LID1_ complex obtained from CG-MD simulations* **Figure supplement 3.** *Snapshots of the different binding states observed for the JAK2-FERM-SH2 – PRLR-ICD_LID1_ complex with the complete structural model of JAK2*

Next, an equilibrated structure of the JAK2-FERM-SH2:PRLR-ICD_K235-E284_ complex was used to build a coarse-grained model, which was then placed near lipid bilayers of different lower leaflet composition. A number of randomly positioned starting orientations were placed ∼7 Å below the lower leaflet (16 orientations for the POPC:POPS:PI(4,5)P_2_ (80:10:10) membrane and eight for the POPC:POPS (70:30) membrane). In addition we included twelve orientations of JAK-FERM-SH2 without the PRLR-ICD_K235-E284_ placed near a POPC:POPS:PI(4,5)P_2_ (80:10:10) membrane. For the JAK2-FERM-SH2:PRLR-ICD_K235-E284_ complex near a 70:30 POPC:POPS membrane, binding to the lower leaflet was observed for only three of the eight systems (***Figure 4 – figure supplement 2A***). In contrast, when PI(4,5)P_2_ was present (10%), rapid binding of the complex to the membrane was observed in all simulations reaching 97% saturation (***Figure 4 – figure supplement 2A***). Both proteins in the complex showed specific clusters of residues with contacts to PI(4,5)P_2_, POPS and POPC, independent of the initial orientations (***Figure 4BC***). The number of contacts formed was higher for the simulations with PI(4,5)P_2_. This suggests that contacts with other components of the membrane occur close to the bound PI(4,5)P_2_. Overall, the PRLR-ICD _K235-E284_ showed a pattern of lipid contacts similar to the simulations of PRLR TMD-ICD_G204-H300_ with the POPC:POPS:PI(4,5)P_2_ (80:10:10) membrane (***Figure 4C***), with residues K_235_GY_237_contacting PI(4,5)P_2_ headgroups for at least 50% of the total contact time and with insertion into the membrane; note that this occurs even though PRLR-ICD_K235-E284_ is not tethered to the membrane via the TMD. Also, like in the PRLR TMD-ICD_G204-H300_ simulations, contacts made by C242 and I243 to POPC were still present. In contrast, contacts by other residues from BOX1 and the KIK motif has lower populations. However, and as expected from the location of the most frequent PRLR-ICD _K235-E284_/lipid contacts, JAK2-FERM-SH2 had more contacts in the F2 lobe of the FERM domain, mainly involving residues I223, L224, R226, K227 and R230, constituting the regions where the N-terminus of PRLR-ICD_K235-E284_ is bound (***Figure 4B***). In the JAK2-FERM-SH2 simulations without PRLR-ICD and near a POPC:POPS:PI(4,5)P_2_ (80:80:10) membrane, we observed that simulations initiated in eleven out of the twelve orientations ended up binding to the membrane (***Figure 4 – figure supplement 2A***). In this case, the overall binding pattern of protein-lipid contacts remained similar.

Previous studies have suggested that the Martini2 forcefield model underestimates cation-π interactions between surface aromatic residues and choline headgroups on the membrane (Khan et al., 2020). However, this may not be applicable to other types of protein-membrane interactions, particularly where negatively charged headgroups are present. Our simulations involving PI(4,5)P_2_) and POPS suggest that interactions between PRLR and the bilayer are primarily driven by positively charged residues in the protein, and that other protein-membrane interactions are secondary or occur between the lipids and residues that surround positively charged residues interacting with a PI(4,5)P_2_ (or POPS) lipid. As a result, cation-π interactions may not be as important for the protein-lipid contact patterns we observed, but could be one explanation as to why we observe less frequent binding in the POPC:POPS systems.

In summary, our simulations showed that binding of JAK2 to the membrane was enhanced by the presence of PI(4,5)P_2_ and that the ICD from PRLR and JAK2 formed a co-structure with PI(4,5)P_2_ maintaining the contacts to the lipids observed for the individual proteins. The presence of PI(4,5)P_2_ was essential for the membrane interactions.

### The complex between JAK2-FERM-SH2 and LID1 shows preferential bound orientations with membranes containing PI(4,5)P_2_

To characterize the membrane-bound modes of the complex in more detail, we took inspiration from Vogel *et al*. (Herzog et al., 2017) and constructed a map that represents the populations of different orientations of the JAK2-PRLR-ICD_K235-E284_ complex relative to the membrane and extracted conformations to represent the most populated orientations as classified into states (***Figure 4D***). For the JAK2-PRLR-ICD_K235-E284_ complex bound to the POPC:POPS:PI(4,5)P_2_ (80:10:10) membrane, States 1–4 (∼31% of the total contact time) showed the complex in an orientation where the N-terminus of PRLR-ICD_K235-E284_ contacted and inserted into the bilayer similarly to what was observed in the PRLR TMD-ICD_G204-H300_ simulations near a POPC:POPS:PI(4,5)P_2_ bilayer. Of these four states, States 1, 2 and 3 had the F2 lobe of the JAK2-FERM domain and the ICJM region of PRLR-ICD_K235-E284_ in contact with the membrane, penetrating below the headgroups, and acting as a pivot over which the protein-complex rotates, leaving the complex to hang as a “Y” from the membrane (See ***Figure 4DE*** and ***MOVIE1***). State 4 on the other hand, while retaining the main contact points, assumed a “flat” orientation with larger sections of the F2 lobe and F1-F2 linker from JAK2-FERM and the entire N-terminal half of PRLR-ICD_K235-E284_ (residues K235-G263) making a substantial number of contacts with the membrane (***Figure 4DF*** and ***MOVIE2***). To examine whether the identified states are compatible with functional states of the full-length kinase, we superimposed representative conformations of States 1 to 6 with that of the full-length JAK2 model obtained from the AlphaFold Protein Structure Database (UNIPROT O60674) (Jumper et al., 2021; Varadi et al., 2022). This procedure revealed that both the Y (States 1, 2 and 3), and Flat (State 4) states keep the other domains of JAK2 oriented towards the cytoplasmic space (***Figure 4 – figure supplement 3AD***), supporting that these states could be functionally relevant. Furthermore, in the context of JAK2 dimerization required for signalling (Ferrao et al., 2018), these states allow for the correct orientation for kinase domain dimerization. The two remaining states (States 5 and 6) showed an inverted orientation in which the main protein-lipid interactions were formed by residues from the F3 lobe of FERM and the SH2 domain, bringing the F2 lobe and the ICJM region of the PRLR-ICD unrealistically far away from the membrane and from the connecting end of the TMD. Thus, States 5 and 6 appear functionally irrelevant, as further demonstrated by the superposition of the full-length AlphaFold model of JAK2 in which the kinase domains would clash with the bilayer (***Figure 4 – figure supplement 3EF***).

### Different membrane co-structures have different exposures relevant to signalling

We analysed the protein-lipid interactions formed by States 2 (Y) and 4 (Flat) in more detail, and despite overall similar contact profiles, some key differences were observed (***Figure 4E–4H***). For the Flat state, an increase in contacts was seen for residues K235 to L260 of PRLR-ICD_K235-E284_ with a pattern similar to the one observed in the PRLR-TMD-ICD_G204-H300_ simulations with PI(4,5)P_2_ present. Dominant PI(4,5)P_2_ contacts were seen for K235–C242, followed by POPC contacts for residues C242–P248, with a second PI(4,5)P_2_ contact peak for K251 and a third around H257. For the Y state, only residues K235–Y237 made substantial contacts with PI(4,5)P_2_ and/or POPC, leaving BOX1 and the KIK motif exposed to the solvent and making contacts with JAK2. For JAK2-FERM-SH2, the main difference between the Y and Flat states was a large decrease in contacts to the F2 lobe in the Flat state accompanied by an increase in contacts in F3 and SH2. Residues in the F2 lobe involved in homodimerization, orientation and activation, including L224, K227, R230 and R234 (Wilmes et al., 2020) were only accessible in the Y state, and not in the Flat state. Thus, we speculate that the Y and Flat states may mimic functionally relevant conformations pertaining to active and inactive states of the signalling complex. Mapping of residues that make contacts with the bilayer in the two states to the conservation maps shows that several positively charged residues of JAK2-FERM-SH2 are largely conserved. Particularly K207, R226, K227 and R228 in F2 with high population contacts with PI(4,5)P_2_ in the Y state are highly conserved. Similar conservation was seen for the positively charged residues K235, K414, K415 and R513, that form contacts with PI(4,5)P_2_ in the Flat state.

Similar simulations were performed near a POPC:POPS 70:30 bilayer, which resulted in only one of eight systems showing stable binding to the bilayer characterized by only one state (***Figure 4 – figure supplement 2A,B***). Here, residues from the F2 lobe of JAK2 form contacts with POPS lipids in a narrow peak containing residues Q219 to R230, while for PRLR-ICD _K235-E284_, residues from the ICJM (K235-C242) and the BOX1 region (C242-P248), make high frequency contacts with both POPS and POPC (***Figure 4 – figure supplement 2DE***). Overall this state is somewhat similar to State 3 observed for the simulations with PI(4,5)P_2_. Similarly, for our simulations of JAK2-FERM-SH2 without PRLR near a bilayer with PI(4,5)P_2,_ 11 out of 12 stable binding conformations revealed three most populated states (E1, E2, E3) (***Figure 4 – figure supplement 2C***). Characterization of these in terms of protein-lipid contact profiles revealed that the positively charged residues in the F2 lobe play an important role in binding to PI(4,5)P_2_ in a similar manner as observed for the Y and Flat states of the complex (***Figure 4 – figure supplement 2F***). Indeed, States 2 and 3 (***Figure4 – figure supplement 2GH***) show remarkable similarities with the Flat and Y state from the simulations of the JAK2-FERM-SH2:PRLR-ICD_K235-E284_ complex near a similar bilayer.

Overall, the simulations highlight preferential binding of both JAK2-FERM-SH2, both alone and in complex with PRLR-ICD_K235-E284_ to PI(4,5)P_2_, and show that the absence of this lipid decreases the level of LID1 binding to the bilayer. Even in the absence of TMD tethering, the most populated bound states recapitulate the binding mode observed for the PRLR-ICD alone. Another key observation is the existence of different states in which different regions of both JAK2 FERM-SH2 domain and LID1 of PRLR are exposed to the solvent or hidden below the bilayer.

### Key residues for membrane interaction control cellular signalling efficiency

From the NMR experiments and MD simulations we identified residues in LID1 that interact with different components of the membrane and/or the FERM-SH2 domain of JAK2. This resulted in four clusters positioned in the ICJM (K235-C242), the BOX1 region (C242–P248), two basic patches of the KxK motif type (K251–K253 and K262–K264) and hydrophobic residues in the region connecting them, respectively. To decipher the specific role of these clusters for PI(4,5)P_2_ interaction, we introduced four sets of mutations in ICD_K235-G313_ and investigated the effect on PI(4,5)P_2_ interaction using NMR spectroscopy. Based on the NMR data and simulations, we focused on the KxK motifs, which would be involved in binding to PI(4,5)P_2_ (***Figure 3A***) and JAK2 (***Figure 4 – sumplement figure 1***), the CIF sequence, indicated to be important to membrane binding (***Figures 2 and 3B***) and four hydrophobic residues, where at least two were seen to be important for JAK2 binding (***Figure 4-supplement figure 1***). We avoided interfering directly with the BOX1 core motif (P245–P250) (Pezet et al., 1997). Thus, the CIF motif (C242–F244) was mutated to GAG (GAG mutant: C242G, I243A, F244G), while the lysines in the KxK motifs (251–253 and 262–264) were all mutated to either glycines (K4G mutant: K251G, K253G, K262G, K264G) or, for charge reversal, to glutamates (K4E mutant: K251E, K253E, K262E, K264E). Finally, four hydrophobic residues (I_252_, F_255_, L_259_ and L_260_) were mutated to glycines (φ4G mutant). ^15^N-PRLR-ICD_K235-G313_ and the four variants were titrated with up to 25× molar excess of C_8_-PI(4,5)P_2,_ keeping the concentration below the CMC. ^1^H-^15^N-HSQC spectra were recorded at each titration point and the changes in chemical shifts and signal intensities were quantified (***Figure 5A***). Similarly to WT, all variants showed negligible chemical shift changes (***Figure 5 – figure supplement 1***) but large peak intensity changes. Decreased peak intensities were observed for all variants in the region of G236-D295, where the changes were largest for the φ4G mutant and K4G, and similar to WT, while smaller effects were seen for the GAG and the K4E variants, suggesting weaker affinities. Together, this indicates that the KxK motifs and the CIF-motif are involved in PI(4,5)P_2_ interaction, as expected from the contacts predicted from simulation and NMR and CD data, yet none of these residues are essential for binding.

**Fig. 5.**
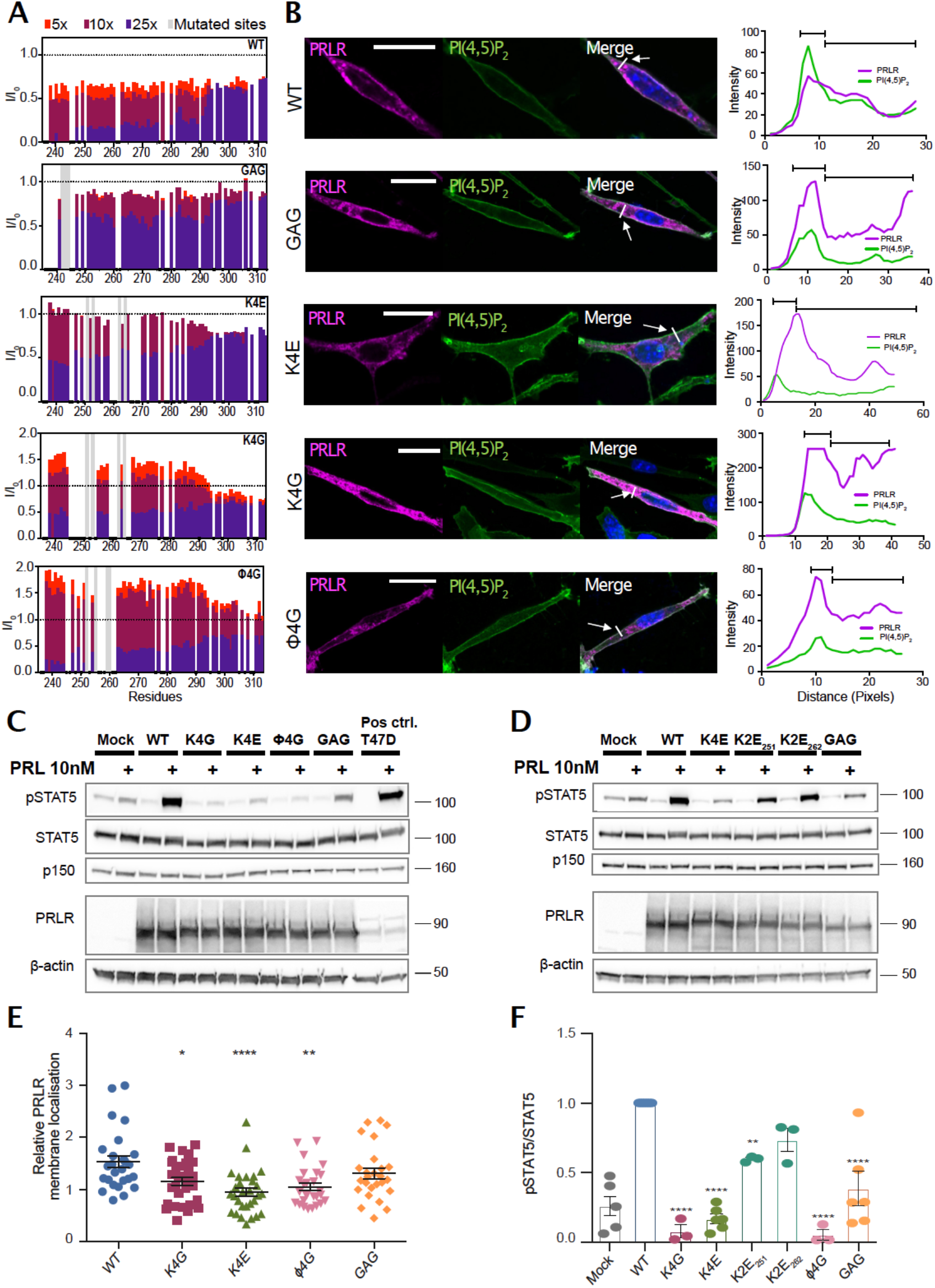
PRLR variants with mutations in lipid interacting residues exhibit decreased PRL-stimulated STAT5 activation in AP1-2PH-PLCδ-GFP cells. **A)** NMR intensity changes of ICD_K235-G313_ WT, K4G, K4E, φ4G and GAG variants upon titration with 5x, 10x and 25x molar excess C_8_-PI(4,5)P_2_ plotted against residue number. **B)** The PRLR variants (WT, K4G, K4E, φ4G, 3GAG) were transiently transfected in AP1 cells stably expressing the 2PH-PLCδ-GFP construct which visualizes the plasma membrane by binding PI(4,5)P_2_. The cells were subsequently analysed by immunofluorescence microscopy, using antibodies against PRLR (magenta) and GFP (green), as well as DAPI (blue) to mark nuclei. To the right, examples of an average line-scan for each PRLR variant is shown. The fluorescence intensity depicted along the white line drawn (arrow) and green fluorescence (plasma membrane) was used to divide the line in a plasma membrane section and intracellular section, and relative membrane localization was calculated as the average fluorescence of PRLR in the membrane section divided by that in the intracellular section. **C, D)** AP1-2PH-PLCδ-GFP cells were transiently transfected with PRLR variants (WT, K4G, K4E, φ4G, 3GAG, K2E_253_, K2E_261_) and incubated overnight followed by serum starvation for 16-17 h and were subsequently incubated with or without 10 nM prolactin for 30 min. The resulting lysates were analysed by western blot for STAT5, pSTAT5 (Y964), PRLR, β-actin and p150 levels. The immunoblots are representative of three biological replicates. **E**) Ratio of plasma membrane localized receptor compared to intracellular receptor, analysed by line-scans as in B. Each point represents an individual cell, and data are based on three independent biological experiments per condition. Graphs show means with SEM error bars. *P<0.05, **P<0.01 and ****P<0.0001. One-way ANOVA compared to WT, unpaired**. F)** Quantification of western blot results shown as pSTAT5 normalized to total STAT5, relative to the WT condition. Graphs show means with SEM error bars. *P<0.05 and **P<0.01. One-way ANOVA compared to WT, unpaired. **Source file 1**: Raw western blots (relating to *figure 5C*) **Source file 2**: Raw western blot (relating to *figure 5D*) **Source file 3**: Data summaries (relating to *figure 5E,F*) **Figure supplement 1**: *Chemical shift perturbations of ICD_K235-G313_ of A) WT, B) K4E, C) GAG, D) K4G and E) ϕ4G variants.* **Figure supplement 2:** *^15^N R_2_ relaxation rates of ICD_K235-G313_ of WT (grey bars), K4G (blue dots) and ϕ4G (red squares) variants.*

We observed dramatic increases in peak intensities for the K4G and φ4G variants in the presence of 5× and 10× molar excess of C_8_-PI(4,5)P_2_ when compared to WT (***Figure 5A***), suggesting changes in the dynamics of the chain. To adress this, we probed the backbone dynamics by acquiring ^15^N *R_2_* relaxation rates of the WT, K4G and φ4G variants in the absence of C_8_-PI(4,5)P_2_ (***Figure 5 – figure supplement 2***). Compared to the WT, no major changes in *R_2_* were observed for two variants. The intensity increase observed for the K4G and φ4G variants during the titration with C_8_-PI(4,5)P_2_ therefore indicates increased backbone dynamics upon binding to C_8_-PI(4,5)P_2_ compared to the WT. Although we cannot explain this observation, it suggests that binding to C_8_-PI(4,5)P_2_ increases the dynamics of the first parts of the chain and thus require higher concentration of C_8_-PI(4,5)P_2_ to fully form the complex, as expected by the lower apparent affinity.

To address the implications of PI(4,5)P_2_ interaction for PRLR membrane localization and downstream signalling, and to enable a potential separation of effect of perturbed membrane localization from direct PI(4,5)P_2_ binding, we introduced the same four sets of mutations into the full-length PRLR. Together with WT PRLR, these were transiently transfected into AP1 mammalian epithelial cells, which were stably transfected with the fluorescent PI(4,5)P_2_ reporter 2PH-PLCδ-GFP. The cells were subjected to fluorescence microscopy analysis of PRLR and the PI(4,5)P_2_ reporter (Fig. 5B) and to western blot analysis of STAT5-activating phosphorylation (***Figures 5C-D, F-G***). None of the mutations fully abolished PRLR membrane localization. Western blot analysis showed that the protein expression levels were similar for WT PRLR and all PRLR variants (***Figure 5C***). However, compared to WT, the K4G, K4E and φ4G variants exhibited a significant reduction in membrane localization as determined by line scan analysis (***Figures 5B,E***). This is in accordance with their predicted JAK2 contacts obtained from the simulations, as JAK2 is known to be important for PRLR trafficking (Huang et al., 2001). The PRL-induced STAT5 activation was significantly decreased in cells expressing either the K4G, K4E or the φ4G variants, whereas STAT5 activation in cells expressing the GAG variant was not significantly different from that of WT expressing cells (***Figures 5C,F***). Decomposing the K4E variant into two individual mutants, in which only one of the two KxK motifs was changed (K2E_251_ and K2E_262_) showed that the reduction in STAT5 activation was attenuated in the variants with the individual mutations, compared to the drastic decrease observed for the K4E mutant (***Figure 5D,G***). Thus, both KxK motifs are important for JAK/STAT activation, which suggests that both PI(4,5)P_2_ and JAK2 binding are important in this regard.

Taken together, these results show that while our data are consistent with the decreased membrane localization contributing to the reduction of STAT5 activation, it is unlikely to account fully for the effect observed for the K4G, K4E or the φ4G variants. Part of this reflects impaired binding to JAK2, known to affect the amount of receptor at the cellular membrane. The MD simulations indicated that only the first KxK motif is involved in lipid interaction while the second KxK motif is involved in JAK2 interaction. Thus, a part of the the reduction in JAK/STAT activation in these variants could arise from a combined effect of abolishing both PI(4,5)P_2_- and JAK2 interaction within the LID1 region, which support the suggestion that co-structure formation between JAK2, PRLR and the membrane is critical for optimal JAK/STAT signalling. However, within this co-structure, the involved residues will likely affect several binding events, and thus separation of function by selective mutations may not be straightforward.

## Discussion

The sequence of the human PRLR has been known for more than 35 years (Boutin et al., 1988). Little attention has, however, been given to the role of membrane composition for PRLR signalling, despite it being placed in the plasma membrane where phosphoinositide levels are highly dynamic and spatially variable, and being linked to cancer with lipid deregulation (Dadhich and Kapoor, 2022). Here we asked if JAK2 and PRLR-ICD share a PI(4,5)P_2_ binding site and if and how the binding to PI(4,5)P_2_ plays a role in the orientation of these proteins with respect to the membrane. Integrating MD simulations with biophysical and cellular experiments has been critical in this endeavor. Our first goal was to identify the residues of LID1 involved, as well as the structure formed—if any—in the protein–lipid complex. Our results suggest that the residues that form the ICJM and BOX1 regions of the ICD interact with the lipids via non-specific hydrophobic interactions that involve penetration of the bilayer below the headgroups. This in turn enables positively charged residues of the _251_KIK_253_ motif to establish ionic interactions with PI(4,5)P_2_ and in doing so the region folds into an extended structure, similar to structures of other cytokine receptors in complex with either JAK1 or TYK2 (Wallweber et al., 2014; Zhang et al., 2016). In turn, PI(4,5)P_2_ lipids accumulate around the TMD and LID1 of PRLR, suggesting a relevant functional role of the interaction.

The results highlight the capacity of the LID1 to establish highly populated and specific interactions with PI(4,5)P_2_ via residues essential for its interaction with JAK2. Therefore, we addressed whether LID1 in complex with the FERM-SH2 domain of JAK2 could engage with the lipid bilayer in the absence of the TMD and with or without PI(4,5)P_2_ lipids. Indeed, PI(4,5)P_2_ was required for binding of the complex to the membrane, as the presence of only POPS in the lower leaflet was not enough to sustain binding despite its negative charge. Remarkably, when PI(4,5)P_2_ was present, we observed specific binding orientations that positioned the ICJM region of the LID1 in the same position as when tethered to the TMD. Furthermore, in the complex, the protein-lipid contact profiles were similar to the one observed for LID1 alone suggesting the PI(4,5)P_2_ binding pattern to be maintained in complex with JAK2. A detailed study of two of the most populated PRLR-bound states of JAK2 revealed a striking difference in orientation and contact pattern with the lipids, that could shed light on functionally relevant states. For example, the most populated state, the Y state, had contacts from the F2 lobe of the JAK2-FERM-SH2 domain and the ICJM of the PRLR, which penetrated the membrane forming hydrophobic interactions with the acyl chains. In this orientation, regions of both JAK2 and the PRLR that have been associated with receptor dimerization and activation for signaling (Ferrao et al., 2018; Wilmes et al., 2020) are exposed to solvent and available for interactions. We note that this orientation has resemblance to that shown in recent cryoEM structures of JAK1 bound to IFNAR1 (Glassman et al., 2022). For the Flat state, we observed a drastic change in the protein-lipid contact profiles for both proteins, but more markedly for LID1. While the main interaction site remains the F2 lobe of JAK2 and the ICJM of the PRLR, the N-terminal residues of the LID1 now lie sandwiched between the membrane and JAK2-FERM-SH2 domain and recapitulates the binding pattern observed from TMD-ICD_F206-H300_ simulations with membranes containing PI(4,5)P_2_. Remarkably, in this Flat state, most of the accessible regions in the Y states are now hidden under the membrane. Thus, we speculate that the Y and Flat states may mimic an available and hidden state, respectively, that could be relevant for regulation of dimerization and activation of signaling. Interestingly, one residue that has been suggested to play a major role in JAK2 membrane association, orientation, dimerization, and activation is L224 (Wilmes et al., 2020). This residue anchors into the membrane only in the Y-state, and not in the Flat-state. Similarly, in simulations where L224 was mutated to glutamate, a change in preferred orientation together with loss of dimerization and JAK2 and STAT5 phosphorylation was observed (Wilmes et al., 2020). This further highlight that the orientation of the PRLR-JAK2 complex relative to the membrane has functional relevance. Importantly, the presence of PI(4,5)P_2_ in the membrane structurally tightens the path from the ECD, folding the ICJM and BOX1 in an extended structure, making transmission of information of hormone binding possible. Thus, signal relay by disordered linker of the PRLR can now be possible through its complex formation with the membrane and JAK2 (***Figure 6***).

**Fig. 6.**
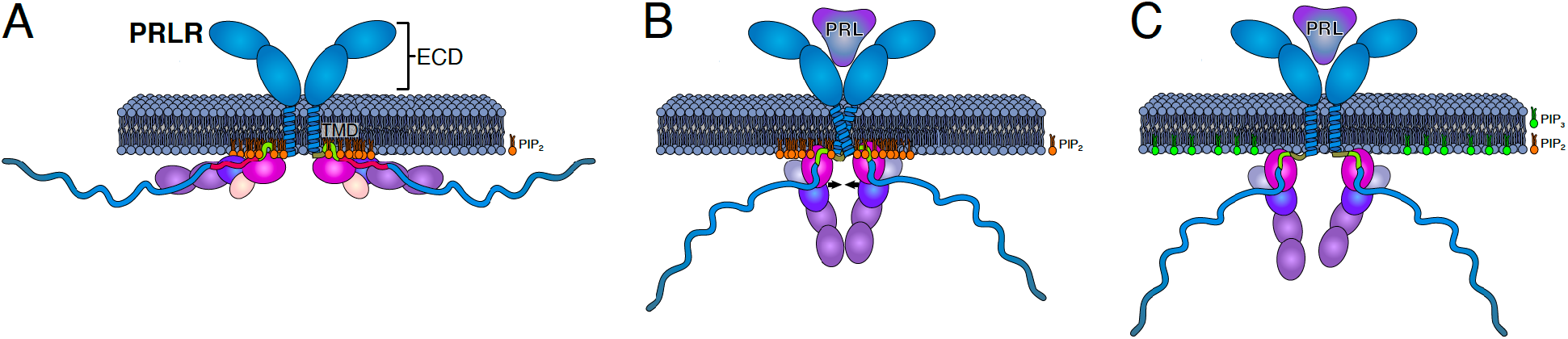
Model of how co-structure formation between JAK2, PRLR and PI4(4,5)P_2_ may contribute to signalling fidelity. The suggested states in signalling would be **A**) the inactive state of the co-structure exemplified by the Flat orientation. **B**) The hormone bound state expemplfied by the co-structure in the Y orientation. **C**) Phosphorylation of PIP(4,5)P_2_ to PI(3,4,5)P_3_ for which the PRLR has no affinity may lead to downregulation and/or termination of signalling. The colour scheme of the proteins is identical to Fig.4.

Lastly, we tested the functional relevance of our observations by mutating residues that appeared significant for PI(4,5)P_2_:PRLR:JAK2 co-structure formation and determining their impact on cellular PRL signaling. Both KxK motifs and the hydrophobic residues connecting them were important for PI(4,5)P_2_ interaction, PRLR membrane localization and cellular JAK2/STAT5 signaling. From the MD simulation it was however evident that not all residues in these motifs were in direct contact with the membrane, further highlighting that co-structure formation between PRLR, JAK2 and the membrane is essential for optimal signal transduction. Another interesting observation was that even though mutating the CIF motif had the largest impact on PI(4,5)P_2_ binding, it had only a limited effect on cellular JAK2/STAT5 signaling. Since the NMR results suggested that the ICJM serves as a primary PI(4,5)P_2_ anchoring point facilitating additional contacts along the chain, this could indicate that a cooperative interaction within the co-structure is needed to control signaling and that PI(4,5)P_2_ interaction is necessary for proper and substantial co-structure formation.

The PI(4,5)P_2_-specific interactions observed point toward a possible regulatory role of PI(4,5)P_2_ in PRLR signaling. Our simulations showed that the membrane embedded TMD-ICD_F206-H300_ was associated with an accumulation of PI(4,5)P_2_ around the TMD. One of the suggested roles of PI(4,5)P_2_ as a regulatory lipid is indeed to form microdomains around proteins and reduce their lateral movement (Trimble and Grinstein, 2015; van den Bogaart et al., 2011). Another possible role can be inferred from previous studies on the EGFR. Evidence suggests a positive feedback loop where inhibition is released upon activation, because PI(4,5)P_2_ is hydrolyzed to DAG and IP_3_ by PLC_y_ (Maeda et al., 2018; McLaughlin et al., 2005). Specifically, we have previously shown that PRLR does not interact with PI(3,4,5)P_3_ (Haxholm et al., 2015). As PI(4,5)P_2_ is phosphorylated by the PI3-kinase to PI(3,4,5)P_3_ during PRLR signaling (Aksamitiene et al., 2011; Yamauchi et al., 1998), this could indicate a way of attenuating signaling. Whether hydrolysis of PI(4,5)P_2_ by PLC_y_ is relevant for PRLR signaling is not known.

## Conclusions

Signal transduction by single-pass receptors through the membrane is still an enigma. In the present work we identify co-structure formation of the disordered LID1 of the PRLR, the membrane constituent PI(4,5)P_2_ and the FERM-SH2 domain of the JAK2, and demonstrate its importance for PRLR signalling. This co-structure has at least two orientations, a Y-shaped state extending from the membrane and a Flat-state with sites hidden in the membrane, the functional roles of which await further elucidation. The co-structure led to accumulation of PI(4,5)P_2_ at the TMD interface and mutation of residues identified to specifically interact with PI(4,5)P_2_ negatively affected PRL-induced STAT5 activation. Facilitated by the co-structure, the disordered ICJM folds into an extended structure, tightening the path from the ECD to the ICD. We suggest that the co-structure formed between receptor, kinase and PI(4,5)P_2_ is critical for signal relay from the extracellular to the intracellular side of the membrane, and that different orientations of the co-structure exist that may represent inactive and active states.

## Materials and Methods

**Expression and purification of TMD** _F206-V240_**, TMD-ICD** _F206-S270_ **and ICD_G236-Q396_** PRLR-ICD_G236-Q396_ was produced as described in (Haxholm et al., 2015), and TMD_F206-V240_ and TMD-ICD_F206-S270_ were produced as described in (Bugge et al., 2015).

### Expression and purification of ICD**_K235-G313_** and variants (K4E, K4G, ϕ4G and GAG)

ICD_K235-G313_ and variants hereof, K4E (K251E, K253E, K262E, K264E), K4G (K251G, K253G, K262G, K264G), ϕ4G (I252G; F255G, L259G, L260G) and GAG (C242G, I243A, F244G) were all produced as follows: Competent BL21(DE3) were transformed using heat shock transformation with pET24a+ plasmids encoding the protein of interest with N-terminal His_6_-SUMO tag. One colony was used to inoculate 10 mL of LB media with 50 μg/mL Kanamycin and incubated overnight at 37 °C at 160 RPM. The overnight culture was used to inoculate 1L M9 minimal media (3 g/l KH_2_PO_4_, 7.5 g/l Na_2_HPO_4_, 5 g/l NaCl, 1 mM MgSO_4_, 4 g/l glucose, 1 g ^15^NH_4_Cl_2_, 1ml M2 trace solution, 50 μg/mL Kanamycin) and grown at 37 °C. At OD600 ∼0.6 recombinant protein expression was induced with 0.1 mM IPTG for 4H at 37 °C. Cells were harvested by centrifugation (5000xg, 20 min, 4°C) and kept at -20 °C until purification. Cells were resuspended in 35 mL Buffer A (10 mM imidazole, 50 mM Tris (pH 8), 150 mM NaCl, 2 mM dithiothreitol (DTT) and lysed with French press at 25 kpsi, followed by centrifugation at 20.000xg, 45 min, 4 °C. The supernatant was applied to 5 mL of pre-equilibrated Ni-NTA beads and incubated for 15 min followed by 50 mL wash with Buffer B 10 mM Imidazole, 50 mM Tris (pH 8), 1M NaCl, 2mM DTT) and 50 mL wash with Buffer A. Protein was eluted with 10 mL Buffer C (250 mM Imidazole, 50 mM Tris (pH 8), 150 mM NaCl, 2 mM DTT). The elution was supplemented with 0.01 mg ULP-1 and dialysed against 1 L of dialysis buffer (50 mM Tris (pH 8), 150 mM NaCl, 1 mM DTT) overnight at 4 °C. The sample was re-applied to the Ni-NTA column and incubated for 15 min. Flow through containing cleaved protein was collected, and the remaining protein was eluted with 10 mL Buffer C. The sample was supplemented with 10 mM DTT before heating at 75 °C for 5 min with gentle rotation of the sample throughout. Sample was transferred directly to ice for 10-15 min incubation followed by centrifugation at 20.000xg, 10 min, 4 °C. The supernatant was concentrated and supplemented with 5 mM betamercaptoenthaonl (b-ME) before application to a HiLoad 16/60 Superdex75 prep grade column equilibrated in 20 mM Na_2_HPO_4_/NaH_2_PO_4_, 150 mM NaCl, 5 mM b-ME (pH 7.3). Fractions containing pure protein were pooled and concentrated

### CD spectroscopy

The peptides covering residues K235-D256 (Pep1) and K253-T280 (Pep2), respectively were purchased from KJ Ross (DK) at 95% purity from HPLC purification. The peptides were dissolved in 10 mM Na_2_HPO4/Na_2_HPO4, pH 7.3 to a final concentration of 40 μM (Pep1) and 25 μM (Pep2) and titrated with TFE or C_8_-PI(4,5)P_2_. The spectra were recorded in a 1mm Quartz cuvette on a Jasco-810 spectropolarimeter purged with 8 l/min N_2_ at 25 °C. A total of 10 accumulations were acquired from 260-190 nm with the following settings: 0.5 nm data pitch, 1 nm band width, response time of 2 seconds, scanning speed of 10 nm/min. A background reference was recorded at identical settings for each sample and subtracted from the relevant spectrum. The spectra were processed by fast Fourier transform filtering and ellipticity converted to mean residual ellipticity ([8]_MRW_)

### NMR spectroscopy

#### *TMD* _F206-V240_ *and TMD-ICD* _F206-S270_

^15^N-labelled or ^13^C,^15^N-labeled TMD-ICD_F206-S270_ was solubilized in molar excess 1,2-dihexanoyl-sn-glycero-3-phosphocholine (DHPC) dissolved in 50 mM NaCl, 20 mM Na_2_HPO_4_/NaH_2_PO4 buffer, pH 7.2. Subsequently, the DHPC embedded TMD-ICD_F206-S270_ was subjected to thorough buffer exchange in a 3 kDa cutoff spinfilter to remove residuals. For reconstitution into POPC SUVs, ^15^N-labeled TMD-ICD_F206-S270_ was solubilized in 300 μL 5:1 methanol:chloroform and mixed with molar excess POPC dissolved in chloroform. The constituents were mixed, followed by evaporation of the organic solvent under a stream of N_2_. When the lipid film appeared dry, it was either left under a stream of N_2_ or placed under vacuum for at least an hour. The resulting proteoliposome film was rehydrated with 1 mL of 50 mM NaCl, 20 mM Na_2_HPO_4_/NaH_2_PO_4_ buffer, pH 7.2, followed by extensive dialysis against the buffer in a 3.5 kDa MWCO dialysis tube at 4 °C. Subsequently, the proteoliposome solution was sonicated in an ultrasonication bath or, if the solution did not clarify, with an UP400S Ultrasonic Processor, in rounds of 2 s with 30 s rest between runs. Finally, the sample was concentrated in a 3 kDa cutoff spinfilter.

All NMR samples of ^15^N-labelled or ^13^C,^15^N-labelled TMD-ICD_F206-S270_ were added 10% (v/v) D_2_O, 2 mM tris(2-carboxyethyl)phosphine (TCEP), 1 mM sodium trimethylsilylpropanesulfonate (DSS), 0.05% (v/v) NaN_3_, and 50 mM NaCl, 20 mM Na_2_HPO_4_/NaH_2_PO4 buffer (pH 7.2) to a final volume of 370 μL followed by pH-adjustment to 7.2 (if needed). All spectra were acquired at 37 °C because the peak intensities of the TMD region decreased at lower temperatures. Free induction decays were transformed and visualized in NMRPipe (F Delaglio et al., 1995) and analysed using the CcpNmr Analysis software (Vranken et al., 2005). Proton chemical shifts were referenced internally to DSS at 0.00 ppm, with heteronuclei referenced by relative gyromagnetic ratios. For assignments of backbone nuclei, heteronuclear NMR spectra of a sample containing 0.5 mM ^13^C,^15^N-labelled TMD-ICD_F206-S270_ in 500 times molar excess DHPC were acquired on a Bruker 750-MHz (^1^H) equipped with a cryoprobe. HNCACB and CBCA(CO)NH spectra were acquired with 32 and 40 of transients, respectively, and 20% non-uniform sampling (Mayzel et al., 2014), and used for manual backbone assignments. SCSs were calculated using random coil chemical shifts from (Kjaergaard et al., 2011) (obtained by supplying primary structure, pH and temperature to the webtool http://www1.bio.ku.dk/english/research/bms/research/sbinlab/groups/mak/randomcoil/script/), which were subtracted from the assigned TMD-ICD _F206-S270_ chemical shifts. The ^1^H,^15^N-HSQC spectrum of 0.4 mM ^15^N-labelled TMD-ICD_F206-S270_ in POPC SUVs (100 times molar excess of POPC) was acquired on a Varian INOVA 750-MHz (^1^H) spectrometer equipped with a room temperature probe. The number of transients was 104.

#### ICD_K235-G313_

ICD_K235-G313_ and the four variants (K4E, K4G, phi4G and GAG) were dialyzed at 4 °C overnight against 20 mM Na_2_HPO_4_/NaH_2_PO_4_ (pH 7.3), 150 mM NaCl. The samples of 50 μM protein were added 1 mM TCEP, 0.25 mM DSS and 10% (v/v) D_2_O and centrifuged at 20.000 xg, 4 °C for 10 min and transferred to 5mm Shigemi BMS-3 tubes. All NMR experiments were recorded at 5 °C on a Bruker Avance III 600 MHz (^1^H) spectrometer equipped with cryogenic probe. Free induction decays were transformed and processed in qMDD (Orekhov and Jaravine, 2011), phased in NMRDraw (Frank Delaglio et al., 1995) and analysed in CcpNMR analysis software (Vranken et al., 2005). Proton chemical shifts were referenced to DSS and nitrogen and carbon to their relative gyromagnetic ratios. ^1^H-^15^N-HSQC experiments were acquired using non-uniform sampling (Mayzel et al., 2014) and recorded on 50 μM ^15^N-ICD_K235-G313_ (or variants) in the absence and presence of 5x, 10x and 25x molar excess of C_8_-PI(4,5)P_2_ (Avanti Lipids 850185).

Transverse ^15^N relaxation rates (*R_2_*) of ICD_K235-G313_ and the two variants, K4G and φ4G, were acquired on Bruker Avance III 600 MHz (^1^H) spectrometer with varying relaxation delays of 0 ms, 33.92 ms, 67.84 ms, 135.68 ms, 169.6 ms, 203.52 ms, 271.36 ms and 339,2 ms, measured in triplicates and peak intensities fitted to single-exponential decays.

### Cell lines and media

AP1-2PH-PLCδ-GFP cells (From J. Snipper; vector from Addgene plasmid #35142) were grown in Minimum Essential Medium Eagle (EMEM, Gibco) containing 10% Fetal Bovine Serum (Sigma Aldrich), 1% penicillin/streptomycin (Sigma), and 1% L-glutamine (Sigma). Stable AP1 clones expressing 2PH-PLCδ-GFP were grown in the same medium, containing 600 µg/mL geneticin (Merck-Millipore) to maintain expression of 2PH-PLCδ-GFP. Cell lines were maintained at 37 °C with 95% humidity and 5% CO_2_ and were passaged by gentle trypsination for a maximum of 15 passages.

### Immunoblotting

Cells were grown to ∼80% confluence in 6-well plates, washed in ice-cold PBS, lysed in boiling lysis buffer (1% SDS, 10 mM Tris–HCl, pH 7.5, with phosphatase inhibitors), boiled for 1 min, sonicated, and centrifuged to clear debris. Identical amounts of protein (12 μg/well) diluted in NuPAGE LDS sample buffer (Novex) with 50% 0.5M DTT were boiled for 5 min, separated on Bio-Rad 10% Tris-Glycine gels, and transferred to nitrocellulose membranes using the Trans-Blot Turbo Transfer system (Bio-Rad). Membranes were stained with Ponceau S to confirm equal loading, blocked for 1 h at 37 °C in blocking buffer (TBST, 5% nonfat dry milk), and incubated with the relevant primary antibodies in blocking buffer overnight at 4°C. After washing in TBST (TBS + 0.1% Tween-20), membranes were incubated with HRP-conjugated secondary antibodies (1:2000, Sigma), washed in TBST, and visualized using ECL reagent (Bio-Rad). Protein bands were quantified by densitometry using ImageJ software, and normalized to those of STAT5 and then to WT.

### Immunofluorescence analysis

For immunofluorescence experiments, cells were grown on 12 mm round glass coverslips to ∼80% confluency and fixed in 2% PFA (30 min at RT). Coverslips were washed three times for 3 x 5 min in PBS, permeabilized for 15 min (0.5% Triton X-100 in TBS), blocked for 30 min (5% BSA in TBST), and incubated with primary antibody in TBST + 1% BSA at RT for 1.5 h. Coverslips were again washed in TBST + 1% BSA, and incubated with AlexaFluor488 and AlexaFluor568 conjugated secondary antibody (1:600 in TBS + 1% BSA) for 1.5 h. Finally, coverslips were incubated with DAPI (1:1000) for 5 min to stain nuclei, washed in TBST, and mounted in N-propyl-gallate mounting medium (2% w/v in PBS/glycerol). Cells were visualized using the Olympus IX83 microscope with a Yokogawa spinning disc confocal unit, using a 60X/1.4 NA oil emersion objective. Image adjustments were carried out using ImageJ software. Line scans were performed using the ColorProfiler ImageJ software plugin.

### Primary antibodies

PRLR (Santa Cruz #SC20992), STAT5 (Santa Cruz #SC835), pSTAT5 (Y964) (Cell Signaling #CS4322), p150 (BD #BD610473), β-actin (Sigma Aldrich #A5441).

### Modelling of simulated proteins

#### PRLR TMD-LID1 on a lipid bilayer

To build a model of the hPRLR-TMD-ICD-LID1region (G204 to H300) we used the MODELLER interface of Chimera (Pettersen et al., 2004; Webb and Sali, 2016). The structure of hPRLR-TMD (PDB 2N7I (Bugge et al., 2016a)) was used as template for the transmembrane helix (in this structure the residue at position 204 (P) was mutated to a G thus, in our model position 204 corresponds to a glycine) and due to the lack of structural templates for the ICD, it was modelled as a disordered coil. This all-atom model was used to build coarse-grained simulation systems where the TMD was embedded in different lipid bilayers composed of POPC in the upper leaflet and either: i) POPC:POPS (70:30), ii) POPC:POPS:PI(4,5)P_2_ (90:5:5) and iii) POPC:POPS:PI(4,5)P_2_ (80:10:10) in the lower leaflet using the CHARMM-GUI martini_maker module (Jo et al., 2008; Qi et al., 2015). The resulting systems were built using the Martini 2.2 forcefield topology and were later adapted to the Martini 3 (version m3.b3.2) (Souza and Marrink, 2020) topology using the martinize2.py tool. For these systems, the PI(4,5)P_2_ parameters were adapted from their Martini2.2 version by changing the names of the beads to the Martini3 naming scheme using as example other available lipids. These Martini3 PI(4,5)P_2_ parameters are available in our github repository (see the Data Avaliability section). Secondary structure restraints from the Martini forcefield were only applied to the TMD and no harmonic bond restraints were defined in the building of these systems.

#### All-atom models of JAK2-FERM-SH2 and its complex with PRLR-ICD_LID1_

To build the JAK2-FERM-SH2 + PRLR-ICD_K235-E284_ complex the following structures where used: JAK1-FERM-SH2 + IFNLR1 (PDB 5L04 (Zhang et al., 2016)), TYK2-FERM-SH2 + IFNAR1 (PDB 4PO6 (Wallweber et al., 2014)) and JAK2-FERM-SH2 (PDB 4Z32 (McNally et al., 2016)). A structural alignment of the three FEMR-SH2 domains was performed with STAMP (Russell and Barton, 1992) using the Multiseq module (Roberts et al., 2006) of VMD (Humphrey et al., 1996). The model of PRLR-ICD_K235-E284_ was generated with the MODELLER interface of Chimera using as template the aligned receptor-ICD regions present on the structures 5L04 and 4PO6. A total 200 models were generated, and the best in terms of its DOPE score (Shen and Sali, 2006) was selected for further use. This resulted in a model of PRLR-ICD_K235-E284_ bound to JAK2-FEMR-SH2. By combining this model with chain A of PDB 4Z32, a structural model of the JAK2-FERM-SH2 + PRLR-ICD_K235-E284_ complex was obtained. All-atom simulation systems were built for JAK2-FERM-SH2 and the JAK2-FERM-SH2+PRLR-ICD_K235-E284_ complex model. The missing residues on the loop of F3 of JAK2-FERM-SH2 were completed using CHARMM-GUI pbd-reader module (Jo et al., 2014, 2008). Hydrogen atoms were automatically added to the protein using the psfgen plugin of VMD (Humphrey et al., 1996). Aspartate, glutamate, lysine, and arginine residues were charged, and histidine residues were neutral. Simulation boxes comprised of solvent and 150 mM NaCl were generated using the CHARMM-GUI solution-builder module (Jo et al., 2008; Lee et al., 2016) using CHARMM36m (Huang et al., 2017) parameters and topologies for the protein and the TIP3P water model for the solvent.

#### Coarsed-grained models of JAK2-FERM-SH2 and its complex with PRLR-ICD_LID1_

To build coarse-grained models of JAK2-FERM-SH2 and complex between JAK2-FERM-SH2 and PRLR-ICD_K235-E284_ complex, a conformation from their respective all-atom MD simulations of the complex was taken after 150 ns (see below). These conformations were used to generate a CG model using the martinize.py script. The Martini 2.2 forcefield (de Jong et al., 2013) was used and intramolecular elastic bonds were defined for JAK2-FERM-SH2 in both systems. To keep the complex formed and to avoid a “collapse” of the disordered PRLR-ICD_LID1_, inter-molecular harmonic bonds were also defined between JAK-FERM-SH2 and PRLR-ICD_K235-E284_ in the complex. In both cases, a force constant of 400 kJ mol^-1^ nm^-2^ and lower and upper elastic bond cut-offs of 5Å and 9Å, respectively were used.

#### Coarse-grained models of JAK2-FERM-SH2 and JAK2-FERM-SH2 + PRLR-ICD_LID1_ near a lipid bilayer

The relaxed CG-model of the JAK2-FERM-SH2 + PRLR-ICD_LID1_ complex or JAK2-FERM-SH2 alone (see below) was placed near (∼ 7Å) pre-equilibrated lipid bilayers with wo different compositions: POPC on the upper leaflet and two differen compositions on the lower leaflet: i) POPC:POPS (70:30), and ii) POPC:POPS:PI(4,5)P_2_ (80:10:10). The systems were solvated with water beads and 150 mM NaCl. A total of 16 initial orientations of the protein were generated by rotating the protein around the x or the y axis (with z being the normal of the membrane).

### Molecular dynamics (MD) simulations

#### Coarse-grained MD simulations

Coarse-grained MD simulations were performed with Gromacs 2016 or 2018 using the Martini 2.2 force field (de Jong et al., 2013) or the open beta version of the Martini 3 (3.b3.2) force field (Souza and Marrink, 2020). For the PRLR-TMD-ICD_K235-L284_ simulations we increased the strength of interactions between protein and water by 10% to avoid excessive compaction of the disordered regions, as has been previously done for IDPs and multi-domain proteins (Kassem et al., 2021; Larsen et al., 2020; Thomasen et al., 2022). Other simulation parameters, common to all the CG simulations performed, were chosen following the recommendations in (de Jong et al., 2016). Briefly, a time step of 20 fs was used, the Verlet cut-off scheme was used considering a buffer tolerance of 0.005 kJ/(mol ps atom). The reaction-field method was used for Coulomb interactions with a cut-off of 11 Å and a relative permittivity of ε_r_ = 15. For van der Waals’ interactions, a cut-off of 11 Å was used. The velocity rescaling thermostat was employed with a reference temperature of T = 310 K, with a coupling constant of τ_T_ = 1 ps. For the equilibrations, the Berendsen barostat was employed (p = 1 bar, τ_p_ = 3 ps), whereas the production runs were performed with a Parrinello-Rahman barostat (p = 1 bar, τ_p_ = 12 ps). A semi-isotropic pressure coupling was used for all the systems that contained a lipid bilayer. For all systems, an initial round of equilibration with decreasing constraints applied to the protein beads and lipid beads was performed following the protocol provided by CHARMM-GUI Martini maker module. For the PRLR-TMD-ICD_K235-L284_ simulations, a total of 11 µs of unconstrained MD were performed of which the first microsecond was considered as equilibration and the last 10 µs as production and used for analysis. For the JAK2-FERM-SH2 and the complex between JAK2-FERM-SH2 in solution, 1 µs of unconstrained simulation was performed. In the case of JAK2-FERM-SH2 or the complex between JAK2-FERM-SH2 and PRLR-TMD-ICD_K235-L284_ near a lipid bilayer, an unconstrained run of 5 µs was performed for each system and the complete trajectory considered for analysis.

#### All-atom molecular dynamics simulations

All-atom MD simulations were performed using GROMACS 2016 and 2018 (Abraham et al., 2015), using the CHARMM36m force field (Huang et al., 2017) for proteins and the TIP3P model for water. The initial system was minimized followed by position restrained simulation in two different phases, NVT and NPT. A 150 ns run of unconstrained NPT equilibration was then performed. The Berendsen thermostat was used for the constrained relaxation runs and the Nose-Hoover thermostat for the production runs. In all cases, the temperature was 310 K. For the NPT simulations, the Berendsen barostat was used during relaxations and the Parinello-Rahman barostat used in unconstrained production runs. In all cases the target pressure was 1 atm. In all the simulations, the Verlet-cutoff scheme was used with a 2 fs timestep. A cutoff of 12 Å with a switching function starting at 10 Å was used for non-bonded interactions along with periodic boundary conditions. The Particle Mesh Ewald method was used to compute long-range electrostatic forces. Hydrogen atoms were constrained using the LINCS (Hess et al., 1997) algorithm.

#### Trajectory analyses

Analysis of the obtained trajectories was performed using VMD plugins, GROMACS analysis tools and in-house prepared tcl and python scripts, available on github (see below). For the characterization of the orientation of the JAK2-FERM-SH2+PRLR-ICD_K235-E284_ complex with respect to the lipid bilayer we used the geographical coordinate system with latitude and longitude devised by Herzog et al. (Herzog et al., 2017). Lipid densities were calculated with the Volmap plugin from VMD considering only the PO4 beads and a 1Å grid. Density plots are shown as an enrichment score with values representing the percentage of enrichment or depletion with respect to the average value on the system as done previously in (Corradi et al., 2018). All molecular renderings were done with VMD (Humphrey et al., 1996). Protein-protein contact maps were calculated using CONAN (Mercadante et al., 2018).

## Acknowledgements

The authors thank the SYNERGY, BRAINSTRUC and REPIN consortia for valuable discussion and Signe A. Sjørup and Katrine Franklin Mark for skilled technical assistance. We are grateful to Dr. Julie Schnipper for generating the AP1-2PH-PLCδ-GFP cells and to Prof. Vincent Goffin for initial discussions regarding the importance of the KxK-motifs in PRLR.

## Additional information

### Funding

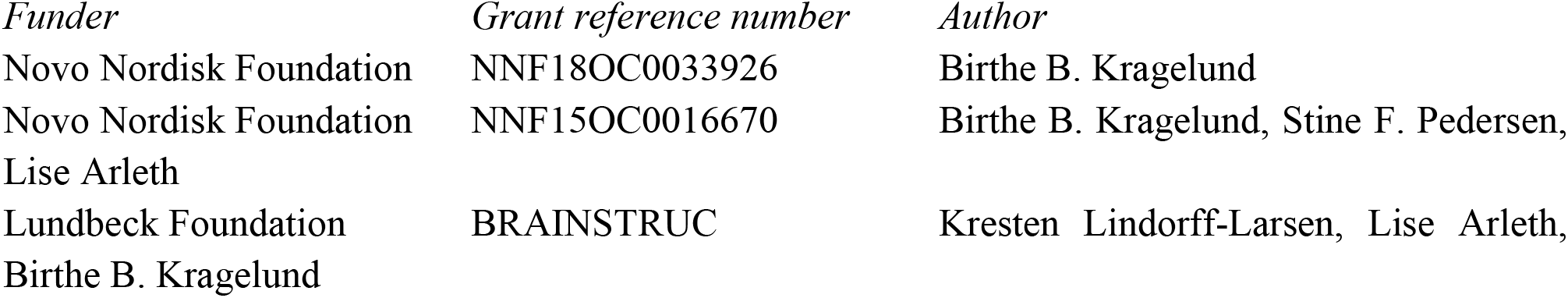

The funders had no role in study design, data collection and interpretation, or the decision to submit the work for publication

### Author contributions

**Raul Araya-Secchi**, Formal analysis, Validation, Investigation, Visualization, Methodology, Writing - original draft; **Katrine Bugge**, Formal analysis, Validation, Investigation, Visualization, Methodology, Writing - review and editing; **Pernille Seiffert,** Investigation, Visualization, Methodology, Writing - original draft, Writing - review and editing; **Amalie Petry**, Investigation; **Gitte W. Haxholm,** Investigation, **Kresten Lindorff-Larsen**, Validation, Supervision, Writing - review and editing, Funding; **Stine F. Pedersen**, Conceptualization, Resources, Formal analysis, Supervision, Funding acquisition, Writing - review and editing;; **Lise Arleth,** Conceptualization, Writing - review and editing; Resources, Formal analysis, Supervision, Funding acquisition; **Birthe B. Kragelund**, Conceptualization, Formal analysis, Validation, Resources, Formal analysis, Visualization, Supervision, Funding acquisition analysis, Writing - original draft, Writing - review and editing;

### Author ORCIDs

Raul Araya-Secchi https://orcid.org/0000-0002-4872-3553

Katrine Bugge https://orcid.org/0000-0002-6286-6243

Pernille Seiffert https://orcid.org/0000-0003-4213-5336

Amalie Petry Gitte W. Haxholm

Kresten Lindorff-Larsen https://orcid.org/0000-0002-4750-6039

Stine Falsig Pedersen https://orcid.org/0000-0002-3044-7714

Lise Arleth https://orcid.org/0000-0002-4694-4299

Birthe B. Kragelund https://orcid.org/0000-0002-7454-1761

## Additional files

### Supplementary files

MDAR checklist

### Data availability

All data needed to evaluate the conclusions in the paper are present in the paper and/or the Supplementary Materials. The MD data and models together with the scripts used in the trajectory analysis are available on Github at https://github.com/Niels-Bohr-Institute-XNS-StructBiophys/PRLRmodel. NMR chemical shifts for the PRLR-ICD_G236-Q396_ are deposited in the BioMagResBank under the accession number 51695.

## Figure supplements and Supplementary files

### Supplementary files not directly related to any main figures

Movie 1: Y State (STATE 2) from the JAK2-FERM-SH2 PRLR-ICDK235-E284 complex simulated near a bilayer containing PI(4,5)P2 : Representative trajectory showing State 2 (Y). Protein and lipids colored as in Fig. 4

Movie 2: FLAT State (STATE 4) from the JAK2-FERM-SH2 PRLR-ICDK235-E284 complex simulated near a bilayer containing PI(4,5)P2 : Representative trajectory showing State 4 (Y). Protein and lipids colored as in Fig. 4

### Figure supplements for main figures

**Figure 2 – figure supplement 1.**
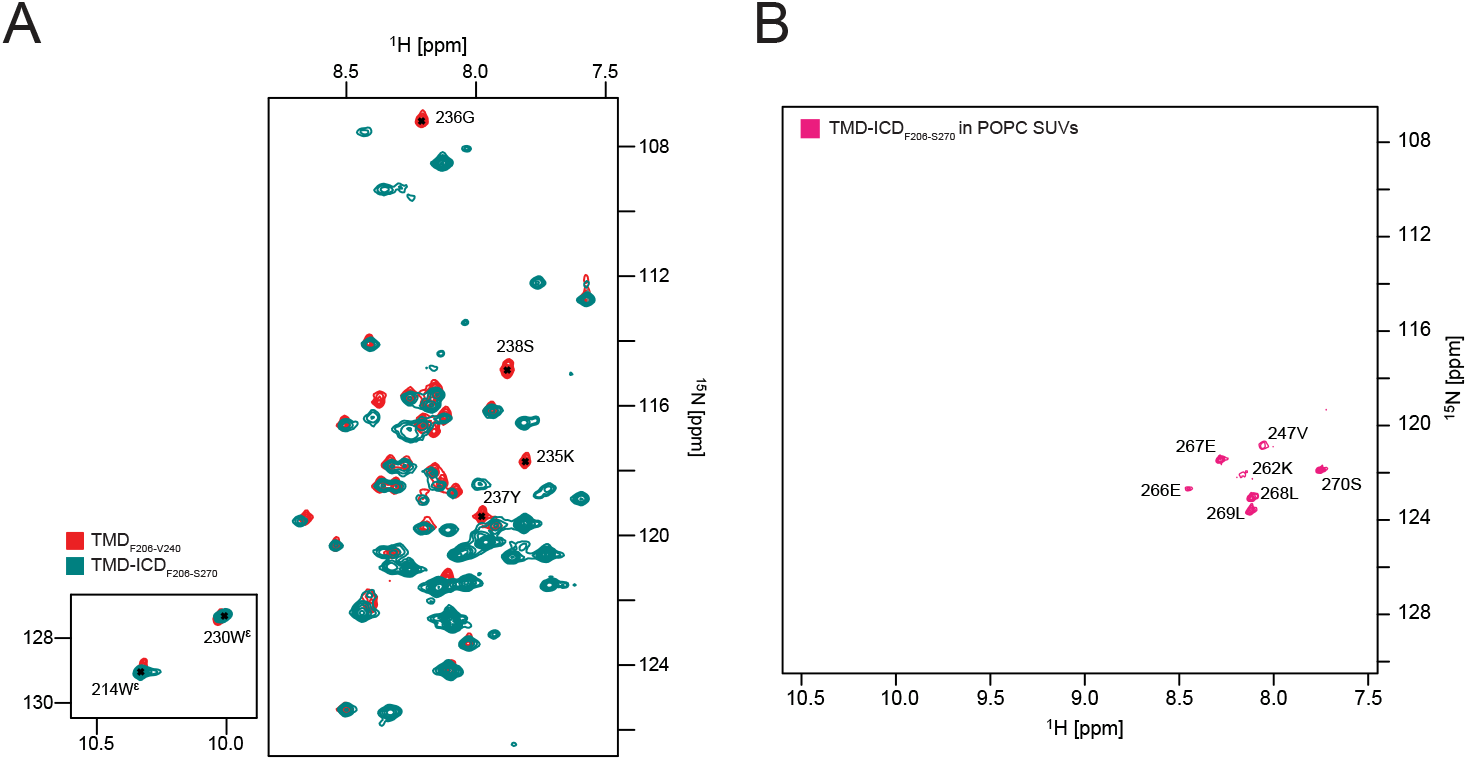
^15^N, ^1^H-HSQC spectra of A) TMDF206-V240 and TMD-ICDF206-S270 in DHPC micelles, and (B) TMD-ICDF206-S270 in POPC SUVs.

**Figure 2 – figure supplement 2.**
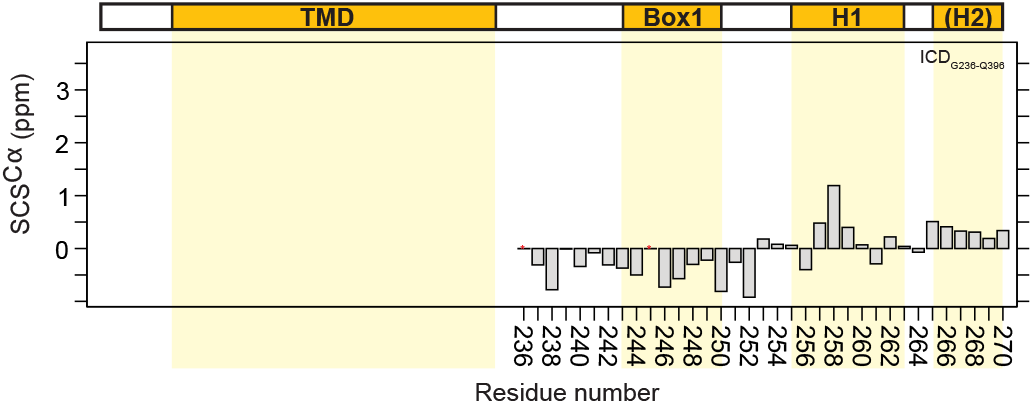
C^α^ secondary chemical shifts of ICDG236-Q396.

**Figure 3 – figure supplement 1.**
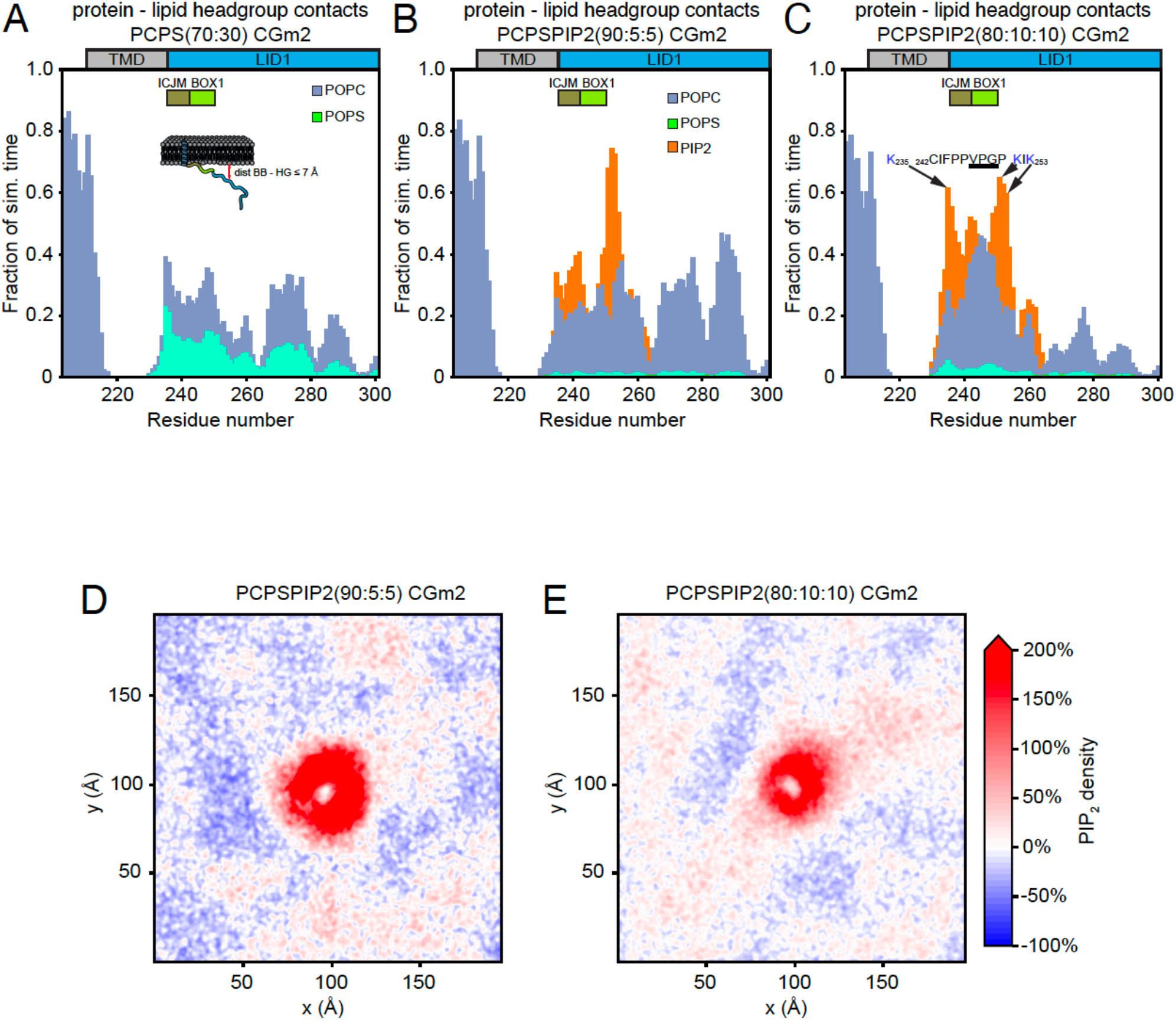
**Protein – lipid interactions of PRLR-ICDLID1 obtained from CG-MD simulations using the martini 2.2 forcefield***. (A-C)* Protein – lipid-headgroups contact histograms from: (A) the PRLR-ICDLID1 POPC:POPS (70:30) CGm2 simulation, (B) PRLR-ICDLID1 POPC:POPS:PI(4,5)P2 (90:5:5) CGm2 simulation and (C) PRLR-ICDLID1 PRLR-ICDLID1POPC:POPS:PI(4,5)P2 (80:10:10) CGm2 simulation.

**Figure 3 – figure supplement 2:**
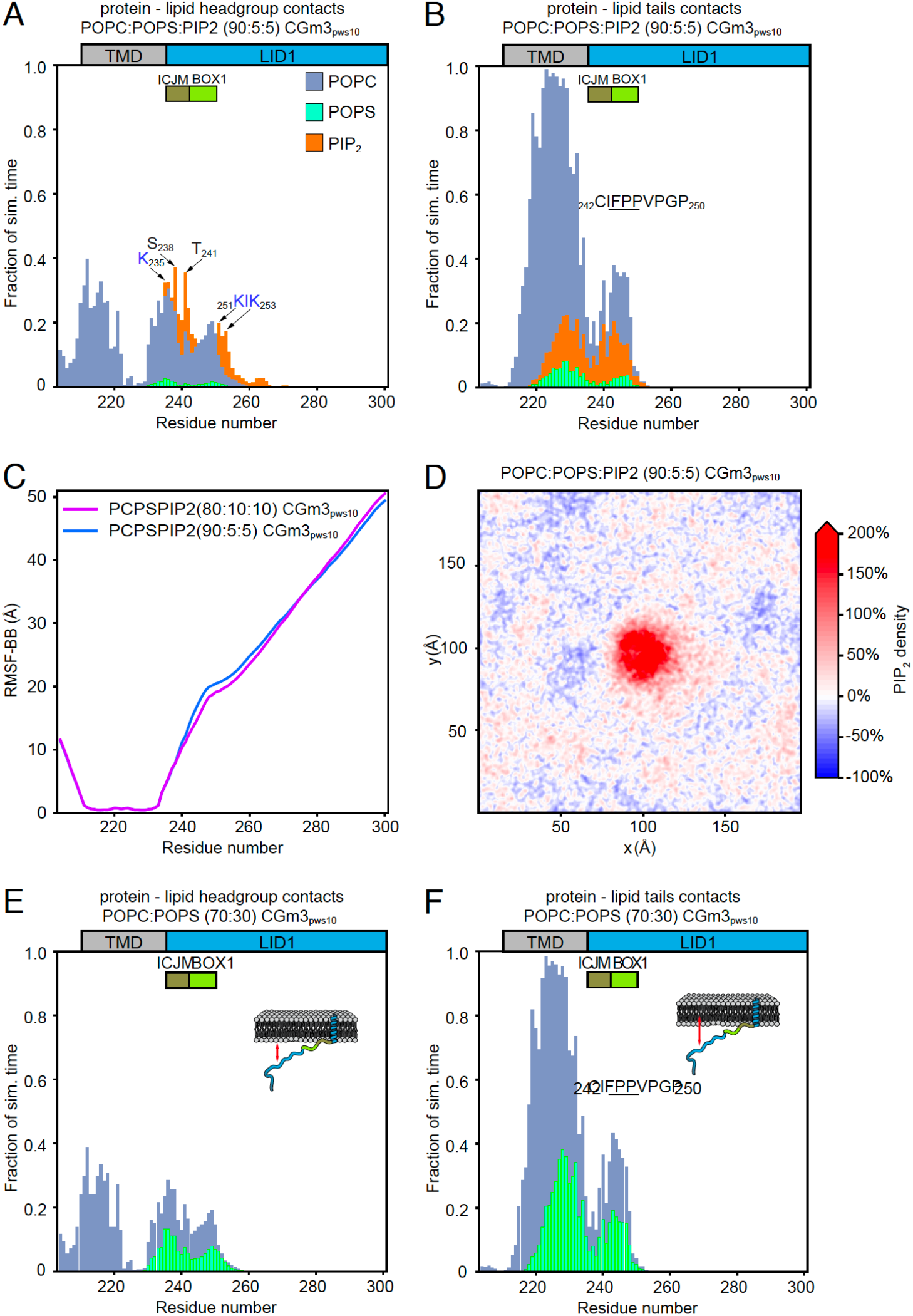
Complementary analysis of protein – lipid interactions of PRLR-ICDLID1 obtained from CG-MD simulations using the martini 3.0b3.2 forcefield. (A-B) Protein – lipids contact histograms from the PRLR-ICDLID1 POPC:POPS:PI(4,5)P2 (90:5:5) CGm3pws10 simulation. (A) Contacts between the protein and lipid headgroups. (B) Contacts between the protein and the acyl chains of the lipids. (C) RMSF of the BB beads obtained from the PRLR-ICDLID1POPC:POPS:PI(4,5)P2 (80:10:10) CGm3pws10 simulation (Magenta line) and the PRLR-ICDLID1 POPC:POPS:PI(4,5)P2 (90:5:5) CGm3pws10 simulation (blue line). (D) Average PI(4,5)P2 density map (*xy*-plane) taken from the PRLR-ICDLID1 + POPC:POPS:PI(4,5)P2(90:5:5) CGm3pws10 simulation. (E-F) Protein – lipids contact histograms from the PRLR-ICDLID1 POPC:POPS (30:70) CGm3pws10 simulation. (E) Contacts between the protein and lipid headgroups. (F) Contacts between the protein and the acyl chains of the lipids. In A-B and E-F protein – lipid contacts are defined as in Figure 3.

**Figure 4 - Figure supplement 1:**
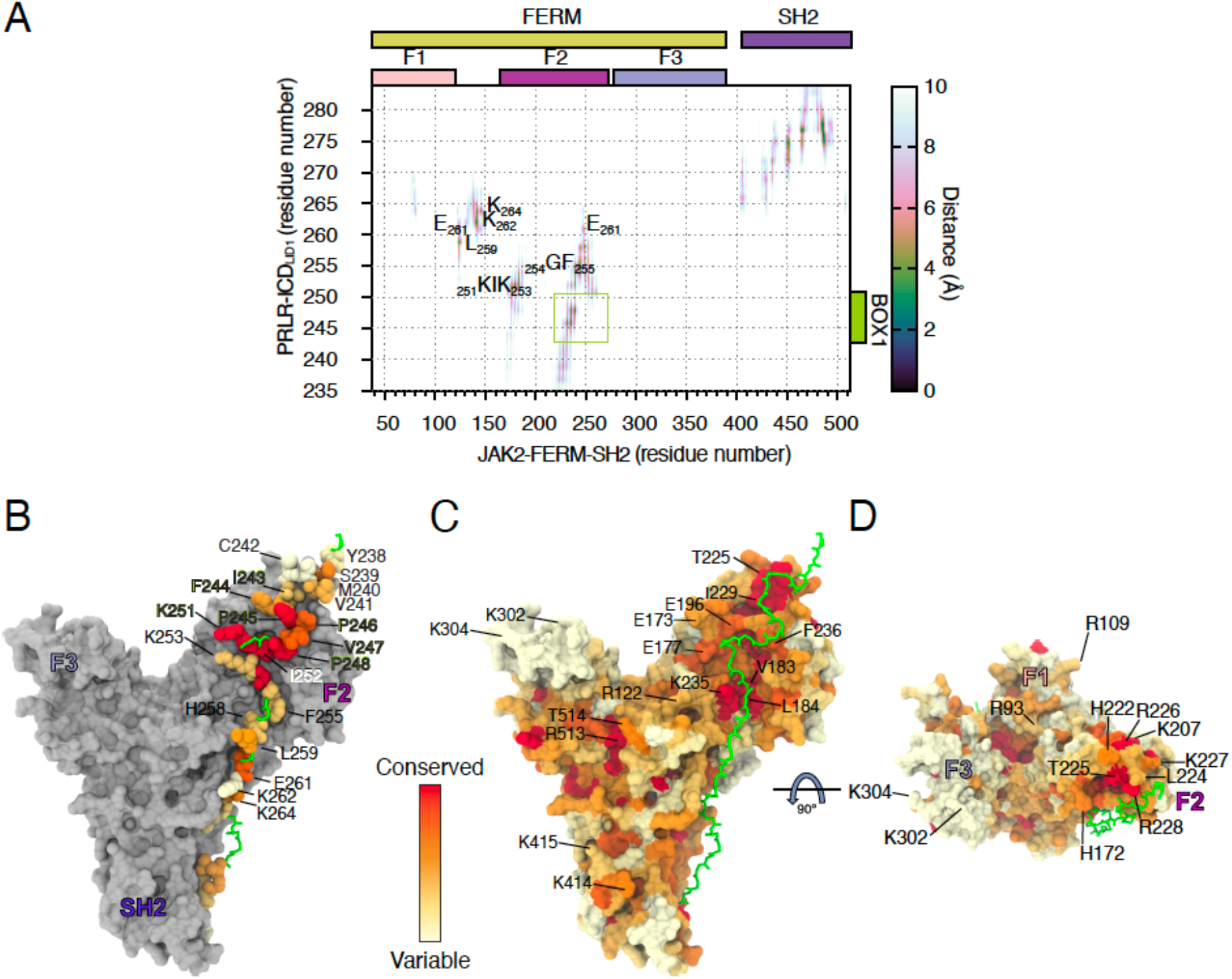
Analysis of the JAK2-FERM-SH2-PRLR-ICDK235-H300 AA-MD simulation. A) Average distance map between JAK2-FERM-SH2 and PRLR-ICDK235-H300 obtained from the all-atom MD simulation. (B) Conservation of residues from PRLR in contact with JAK2-FERM-SH2 (see A). Residues with green label correspond to BOX1. (C-D) Conservation JAK2-FERM-SH2 oriented as state 2 (see Fig. 4E). Labeled residues correspond to residues that contact PRLR (see A) or that contact the lipids on states 2 and 4 (see Fig. 4E-H).

**Figure 4 - Figure Supplement 2.**
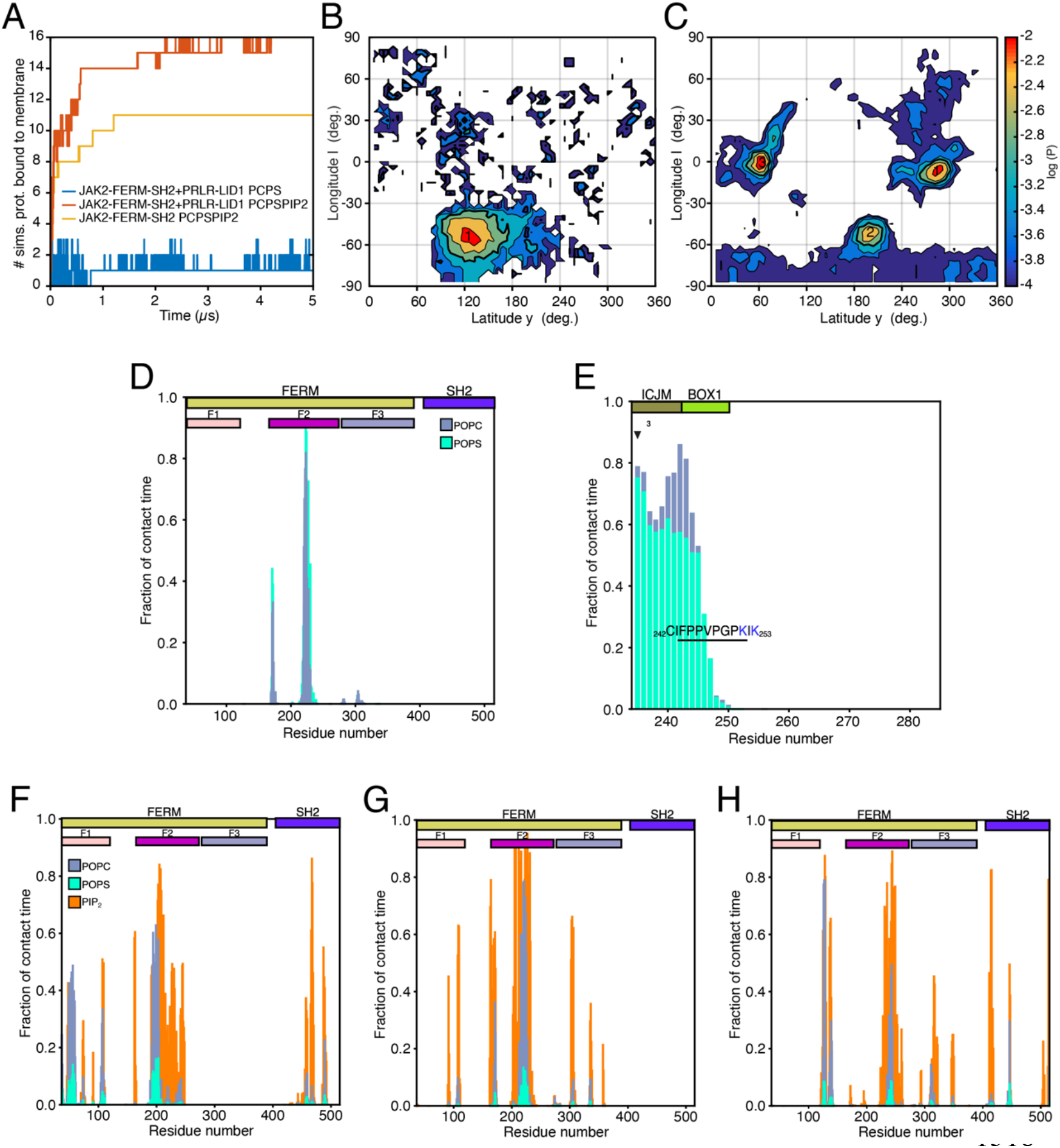
Complementary analysis of Protein – lipid interactions of the JAK2-FERM-SH2 PRLR-ICDK235-H300 complex obtained from CG-MD simulations. (A) Number of simulations where protein is bound to the lower-leaf of the bilayer for the simulations containing: the JAK2-FERM-SH2 + PRLR-ICDK235-H300 complex near a POPC:POPS(70:30) bilayer (blue line) and the JAK2-FERM-SH2 + PRLR-ICDK235-H300 complex (red line) and the apo JAK2-FERM-SH2 (orange line)near a POPC:POPS:PI(4,5)P2 (80:10:10) bilayer. (B-C) Distribution of the orientations adopted by (B) the JAK2-FERM-SH2 + PRLR-ICDK235-H300 complex when bound to lipids taken from the simulations with POPC:POPS(70:30) in the lower-leaflet and (C) the apo JAK2-FERM-SH2 when bound to lipids taken from the simulations with POPC:POPS:PI(4,5)P2 (80:10:10) in the lower-leaflet. (D-E) Representative protein-lipid contact histograms for State 1 observed for the JAK2-FERM-SH2 + PRLR-ICDK235-H300 complex near a POPC:POPS(70:30) bilayer. (F-H) Representative protein-lipid contact histograms for States 1, 2 and 3 observed for the apo JAK2-FERM-SH2 near a POPC:POPS:PI(4,5)P2 (80:10:10) bilayer.

**Figure 4 - Figure Supplement 3.**
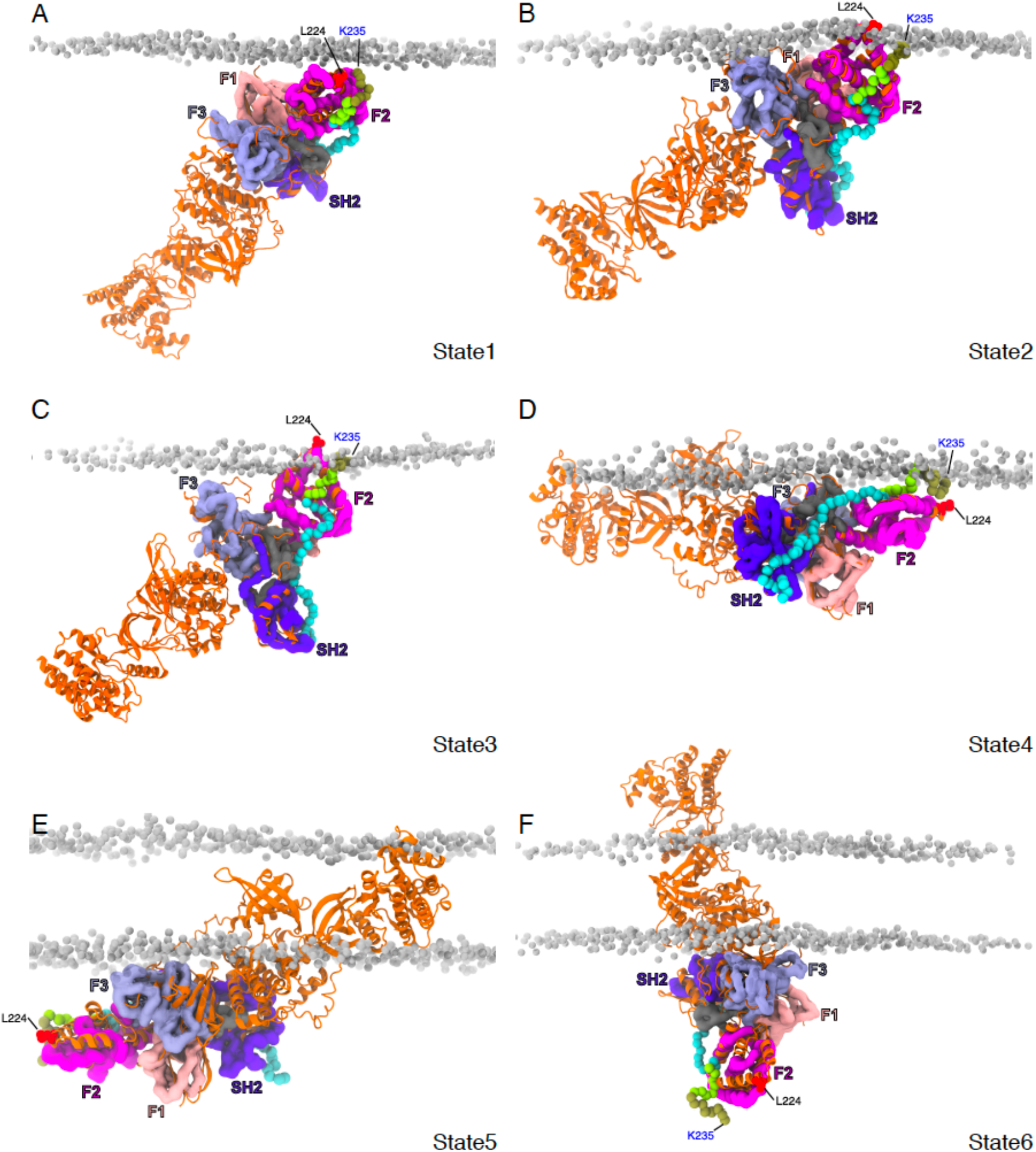
Snapshots of the different binding states observed for the JAK2-FERM-SH2 – PRLR-ICDK235-H300 complex with the complete structural model of JAK2. (obtained from AF2-EBI database). In each panel the CG JAK2-FERM-SH2 – PRLR-ICDK235-H300 complex is depicted and colored as in Fig. 4. The full-length JAK2 model is shown in cartoon representation colored orange. near a POPC:POPS(70:30) bilayer. (F-H) Representative protein-lipid contact histograms for States 1, 2 and 3 observed for the apo JAK2-FERM-SH2 near a POPC:POPS:PI(4,5)P2 (80:10:10) bilayer.

**Figure 5 – Figure supplement 1:**
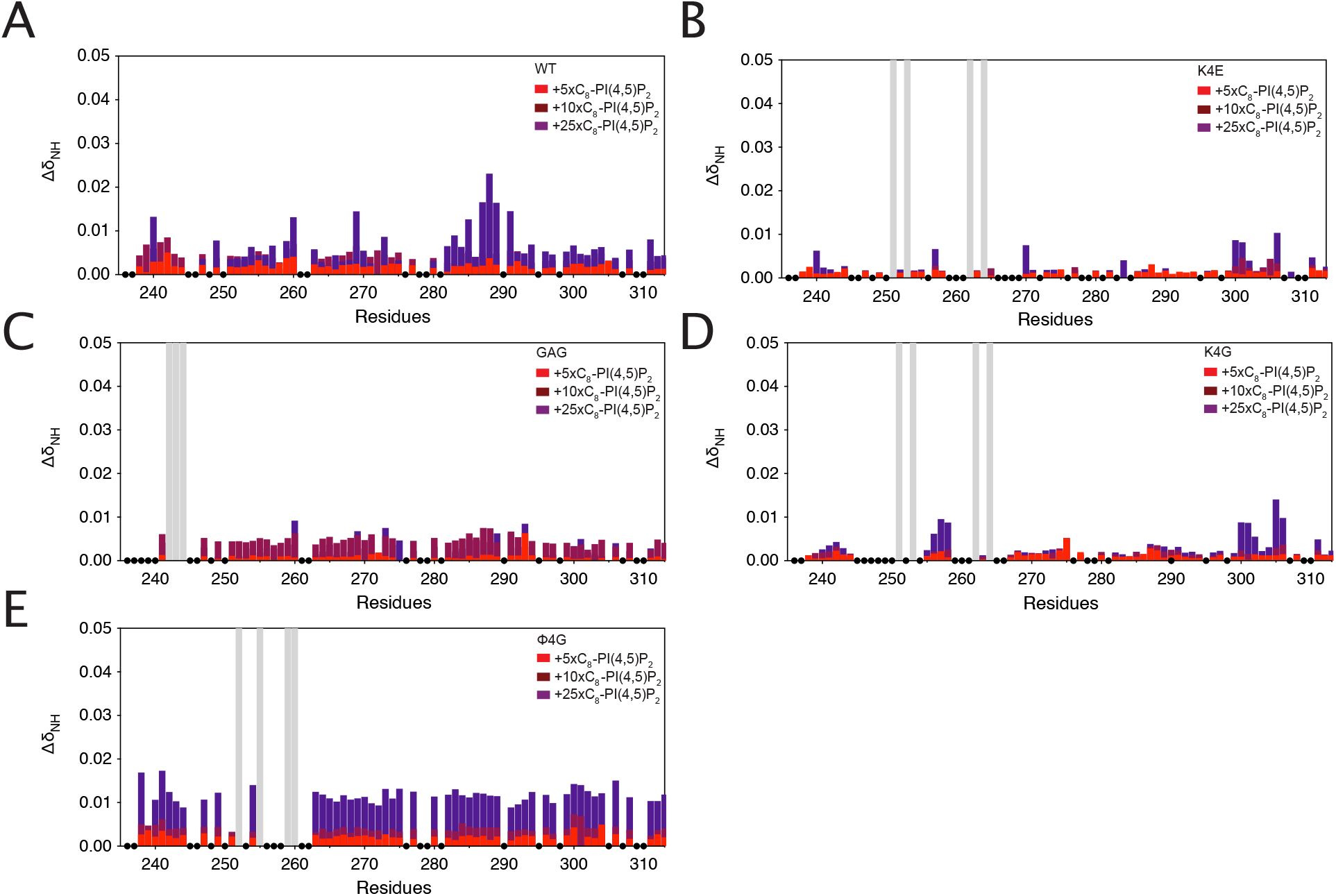
Chemical shift perturbations of ICDK235-G313 of A) WT B) K4E, C) GAG, D) K4G and E) ϕ4G variants.

**Figure 5 – Figure supplement 2:**
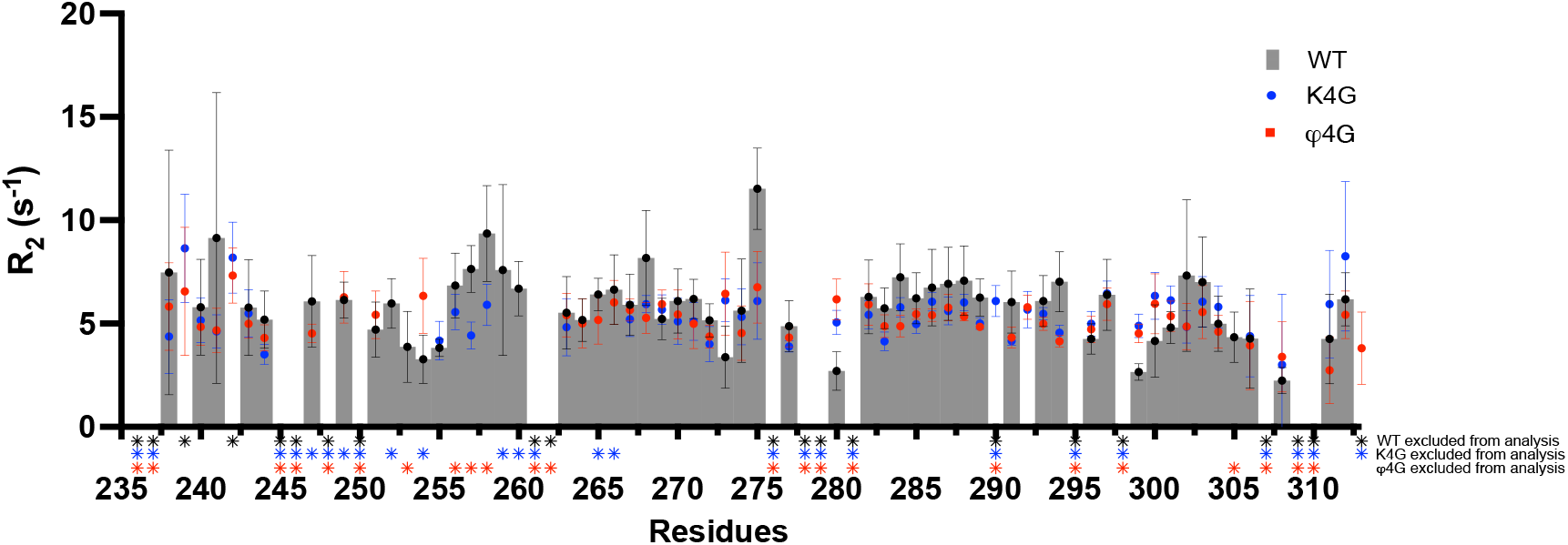
^15^N *R2* relaxation rates of ICDK235-G313 of WT (grey bars), K4G (blue dots) and ϕ4G (red squares) variants. **Stars** indicate signals excluded from the analysis either because it is a proline, signals are overlapping or a poor exponential fit.

